# The balanced lethal system in *Triturus* newts originated in an instantaneous speciation event

**DOI:** 10.1101/2024.10.29.620207

**Authors:** James France, Manon Chantal de Visser, Jan W. Arntzen, Wiesław Babik, Milena Cvijanović, Ana Ivanović, Jeramiah Smith, Tijana Vučić, Ben Wielstra

## Abstract

*Triturus* newts are afflicted by a balanced lethal system causing the spontaneous death of half of their offspring. How could such a maladaptive trait evolve? We construct genetic maps for *Triturus* and its sister genus *Lissotriton*, identifying genes involved in the balanced lethal system. *Triturus* chromosome 1 has diverged into two versions, which each carry a single massive deletion that is compensated for by duplication of the same region in the alternate version – indicating that the balanced lethal system arose instantaneously, in a single macromutation. Simulations show that, counterintuitively, the deleterious nature of the rearranged chromosomes protects them against competition with the ancestral arrangement via reproductive isolation. We conclude that the origin of the *Triturus* balanced lethal system also led to instantaneous speciation.

## Introduction

Evolution is a seemingly simple process that continually produces complex and counterintuitive outcomes, occasionally appearing to contravene even principles as fundamental as natural selection. Investigation of these paradoxical phenomena often leads to a deeper understanding of evolutionary processes. An extreme example of life apparently defying natural selection is found in crested and marbled newts (the genus *Triturus*). In these species, 50% of all embryos spontaneously die before hatching due to a phenomenon known as a balanced lethal system (*1–3*).

The premature death of these embryos is linked to *Triturus’* chromosome 1, which occurs in two distinct versions, termed 1A and 1B. Only heterokaryotypic individuals, possessing both versions of the chromosome, are viable (*2, 4*). Because chromosome 1 is inherited in a Mendelian fashion, and all adults must possess one copy of each version, half of all offspring will inherit two copies of the same version (Fig. 1). The death of these homorokaryotypic embryos implies that each version of chromosome 1 must carry unique mutations that are recessively lethal. Normally these mutations would be selected against and eventually go extinct. However, because the alternative version of the chromosome also has its own lethal mutations, the selective forces are balanced and both chromosomes are maintained in the population, at the cost of half of the offspring, every generation (*3, 5*). Given this extreme fitness penalty, a naturally occurring balanced lethal system seems almost impossible. Yet, balanced lethal systems have independently evolved in widely divergent taxa, from plants to insects. (*6–8*).

**Fig. 1:**
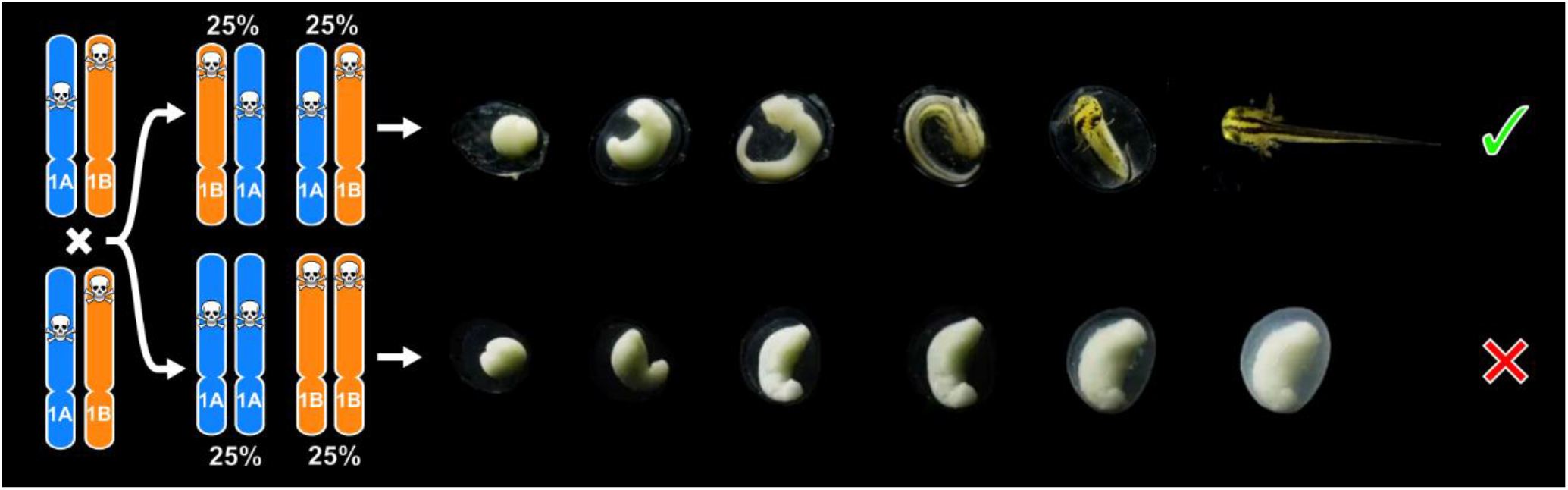
The balanced lethal system on *Triturus* chromosome 1. All adult crested and marbled newts carry two distinct versions of their largest chromosome (termed 1A and 1B, shown in blue and orange). Each version carries unique, lethally recessive alleles. Mendelian segregation results in 50% of offspring inheriting two copies of the same version, but these embryos never hatch, instead suffering arrested embryogenesis and eventual death.

To maintain a balanced lethal system, the chromosomes involved must be shielded from the shuffling effect of recombination. In *Triturus* the long arms of chromosome 1A and 1B form a non-recombining section where no chiasmata are observed (*9*). Non-recombing regions may be created by chromosomal rearrangements, such as inversions, and are associated with phenomena such as sex chromosomes and supergenes (where multiple genes coding for diverse traits are consistently inherited together) (*10*). Many non-recombining regions are lethal when homozygous, for example the supergenes found in ruffs and fire ants (*11, 12*). Consequently it has been proposed that a balanced lethal system may evolve from a degenerate sex chromosome (*13*) or supergene (*14*) system, or may be generated instantaneously by a chromosomal rearrangement (*4*). However, due to the unwieldy size of the *Triturus* genome (ca. 30 Gbp) (*2*), no studies have investigated these hypotheses at a molecular level.

## Genomic architecture

We aim to identify the genomic architecture inhibiting recombination in *Triturus* chromosome 1. We construct a high-density linkage map consisting of 4,226 nuclear DNA markers sequenced in 206 full-sibling offspring of a *T. ivanbureschi* × *T. macedonicus* F_2_ cross. These offspring include approximately equal numbers of viable (heterokaryotypic, designated AB) and arrested (homokaryotypic, designated AA or BB) embryos. As a proxy for the ancestral state of the *Triturus* genome, we construct an analogous map for its sister genus *Lissotriton* (*15*), which diverged approximately 35 million years ago (*16*), before the evolution of the balanced lethal system. The *Lissotriton* map consists of 3,693 markers, of which 2,551 are also located in the *Triturus* map (Figs. S1, S2, Tables S1, S2).

We compare these linkage maps to each other, and to the chromosome-scale genome assemblies of the Iberian ribbed newt, *Pleurodeles waltl*, (*17*) and axolotl, *Ambystoma mexicanum* (*18, 19*). Synteny between *Triturus* and *Lissotriton* is highly conserved, with 98% of genes placed in orthologous chromosomes (Figs. 2A, S3-5). This confirms that the rearrangement that led to the balanced lethal system is restricted to *Triturus’* chromosome 1. Despite over 60 million years of divergence since their last common ancestor (*16, 20*), synteny is also highly conserved between *Triturus* and *Pleurodeles*. Beyond the family Salamandridae, we show that *Triturus’* chromosome 1 is homologous to a fusion between *A. mexicanum* chromosomes 8 and 13.

**Fig. 2:**
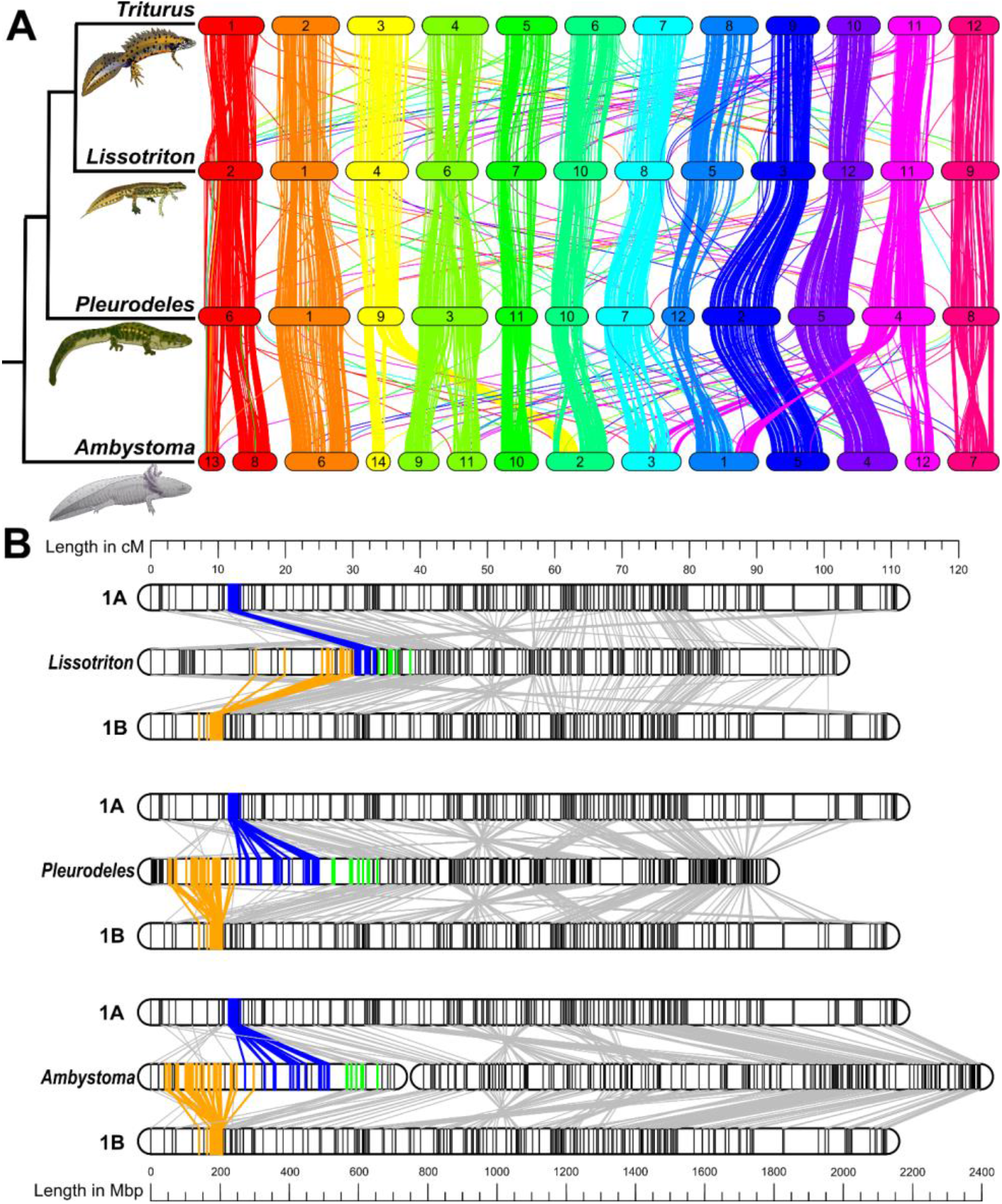
Comparative genomics reveals the *Triturus* balanced lethal system. We construct linkage maps for *Triturus* and *Lissotriton* based on target capture data for ca. 4k coding sequences and incorporate data from published whole genome assemblies of *Pleurodeles* and *Ambystoma* (*17–19*). (**A**) At the whole genome level, we observe a striking conservation of synteny within the three newt genera (*Triturus, Lissotriton* and *Pleurodeles*), with all chromosomes showing one-to-one homology and little variation in gene order. When compared with *Ambystoma* we observe some fusions and translocations (sup. figs. S3-4) (**B**) Linkage groups 1A and 1B of the *Triturus* map compared to the homologous linkage group 2 of *Lissotriton*, chromosome 6 of *Pleurodeles*, and chromosomes 8 and 13 of *Ambystoma*. Genes present only on chromosome 1A are highlighted in blue, those present only on 1B in orange. In all three comparisons, the homologs of these genes present as two distinct adjacent blocks – which have each been entirely deleted from one of the two versions of chromosome 1 in *Triturus*. Genes shown in green form a third block, showing species-specific chromosome 1 related variation (*21*).

Rather than SNPs specific to chromosome 1A or 1B, the variation between genotypes is primarily characterized by genes that are completely absent in one of the two categories of arrested embryos. We identify 30 markers which consistently fail to yield reads in embryos of genotype BB, and 35 are similarly missing in genotype AA – we designate these A-linked and B-linked genes respectively. These two sets of genes are almost identical to those independently discovered by de Visser et al. (*21*), who also report that both sets of genes show highly consistent presence/absence variation across *Triturus* species, indicating that the balanced lethal system attained its modern form before the radiation of the genus.

Remarkably, when the orthologs of the A- and B-linked genes are highlighted in the genomes of *Lissotriton, Pleurodeles* and *Ambystoma*, they are observed to form two distinct, adjacent, but non-overlapping blocks, each corresponding to genes present only in one form of chromosome 1 (Fig. 2B). This shows that the balanced lethal system results from a pair of large deletions – the sizes of the orthologous blocks in the *Pleurodeles* genome are 227 and 189 Mbp for the A- and B-linked genes. As each of the deleted blocks is only present on one form of chromosome 1, recombination between the two forms is impossible in this region.

Cytological studies report that the non-recombining region occupies approximately half of *Triturus* chromosome 1, with an estimated size of 1.3 Gbp (*2*). In contrast, we observe that a large majority of genes on *Triturus* linkage group 1 are fully recombining. This discrepancy does not appear to be an artifact of linkage map construction, as all regions of the homologous chromosomes of other genera are accounted for within the *Triturus* map. A possible explanation is accumulation of repetitive DNA within the non-recombining region, which is unable to purge these sequences (*22, 23*).

### Twin deletions or unequal exchange?

While deletions of the magnitude we observe in *Triturus* would almost certainly be lethal when homozygous, they should also be expected to be deleterious in heterozygous individuals. Given the number of genes involved, it is likely that at least some are haploinsufficient, meaning that a single copy would be insufficient to produce a normal phenotype – for reference, approximately 10% of human genes exhibit haploinsufficiency (*24*). These dosage effects could be compensated for if each region deleted from one version of the chromosome was duplicated on the opposite version, and *vice versa*. As Sessions et al. (*22*) suggested, an unequal exchange between sister chromosomes (or possibly mitotic recombination between homologous chromosomes) would result in exactly this configuration, with the A-linked genes on one chromosome swapped for the B-linked genes on the other (Fig. 3A).

**Fig. 3:**
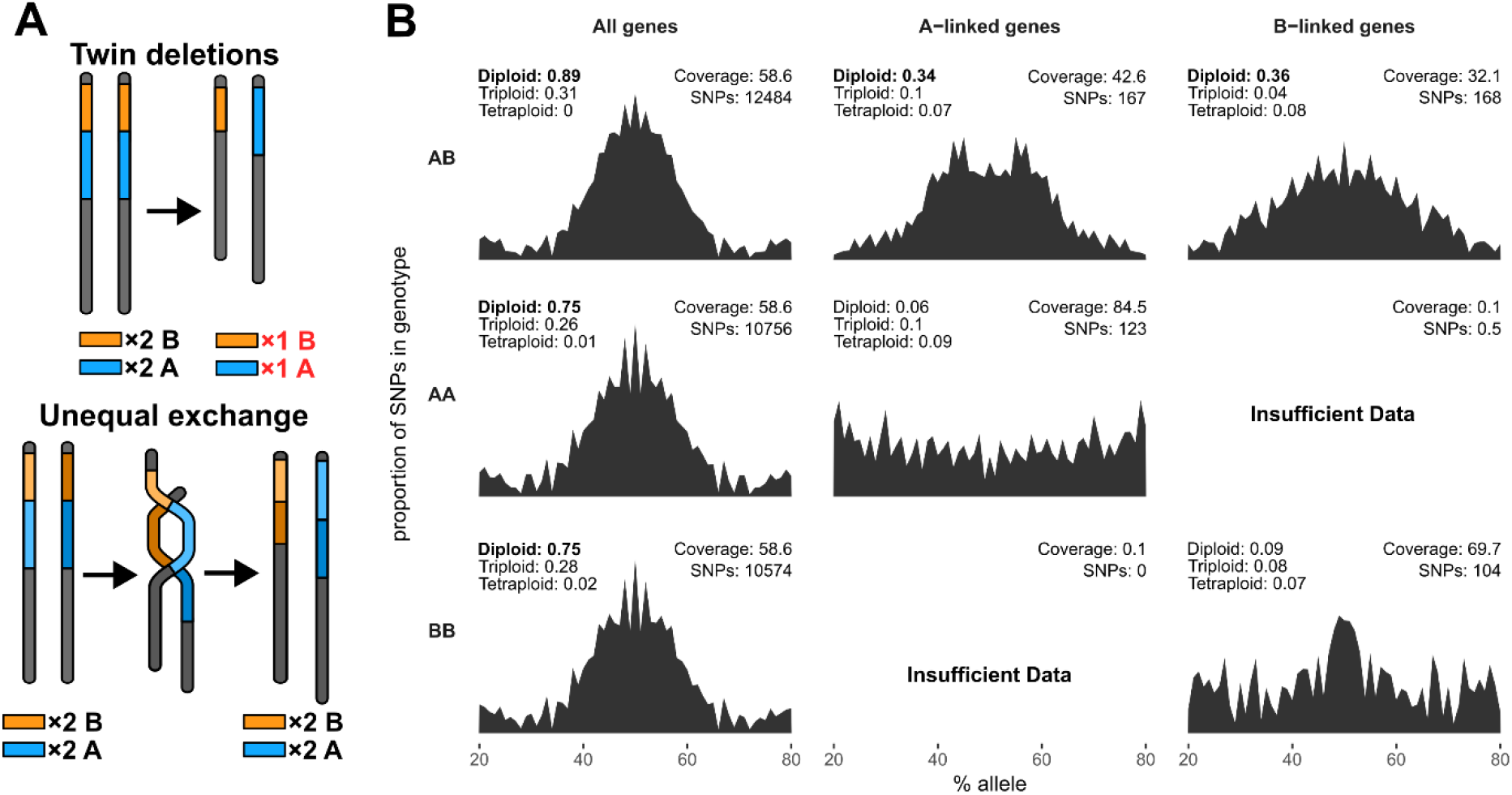
Twin deletions or unequal exchange? (**A**) Two mechanisms may explain our observation that a block of genes is missing from each version of *Triturus* chromosome 1. If each block has been subject to a simple deletion (top) then the genes involved will be left as single copy. Alternatively, If the deletions are part of an unequal exchange (bottom) they will be compensated for by duplication of the same stretch of DNA on the opposite chromosome, leaving two copies. (**B**) We test copy number by measuring allele ratio distribution across three categories of genes in 30 F1 *T. ivanbureschi* × *T. macedonicus* F1 embryos (*25*), split evenly between three genotypes (AB, AA and BB). Coefficient of determination (R2) values comparing the observed results to idealized distributions for different copy numbers are displayed - bold where there is significant agreement. Using the entire set of 7,139 target sequences as a control (“All genes”) we observe a clear diploid distribution, with SNPs tending towards an allele ratio of 50%. Crucially, we also observe this diploid distribution in A- and B-linked genes in the AB embryos despite these individuals only having one copy each of chromosome 1A and 1B, indicating that these genes are indeed duplicated.

If the deleted regions in the *Triturus* balanced lethal system are compensated for by reciprocal duplications, then there will be two copies of A- and B-linked genes in healthy (AB) *Triturus* embryos, even though they only possess a single copy of chromosome 1A and 1B. We test this prediction by analyzing the allele ratio of SNPs in A- and B-linked genes (Fig. 3B) using data from a set of 30 F_1_ *T. ivanbureschi* × *T. macedonicus* (10 for each genotype) (*25*). If there is only a single copy, then an allele can be present in either 0 or 100% of the reads covering each SNP locus within a sample. Instead, we observe that A- and B-linked genes in healthy (AB) embryos possess SNPs where the percentage of reads carrying each allele tends towards 50%, strongly supporting diploidy for these genes and indicating that the A- and B-linked regions have indeed been duplicated.

In arrested embryos of genotype AA, there are two copies of chromosome 1A, so unless the duplication has occurred, we would expect a diploid distribution of SNP alleles in A-linked genes. However, we observe a chaotic distribution, with no significant support for diploidy. Considering the results of the AB embryos, we suspect this is a tetraploid distribution of allele ratios (with peaks at 25, 50 and 75%) that has been obscured by noise. As there are no copies of B-linked genes in the AA samples, we are unable to calculate allele ratios (Tables S3-5). The equivalent allele ratio distributions are found in reverse for the arrested embryos of genotype BB. These results match with the predictions of the reciprocal duplication. Accordingly, we conclude that the *Triturus* balanced lethal system arose in a single step, as the result of an unequal exchange.

### Evolutionary scenario

Our investigation of the *Triturus* genome reveals the architecture of the balanced lethal system, but how could such a deleterious mutation become fixed in the population? Previous work simulating the evolution of balanced lethal systems assume gradualistic scenarios. Grossen et al. (*13*) propose that *Triturus*’ chromosomes 1A and 1B were originally two different lineages of a Y-chromosome, which evolved into the current system due to extinction of the X-chromosome. Berdan et al. (*14*) model the origin of a balanced lethal system as an effect of a self-reinforcing heterozygote advantage, resulting in the gradual accumulation of lethal alleles. However, at least for *Triturus*, these models are incompatible with the genomic evidence, which supports instantaneous evolution.

This result seems paradoxical. As Grossen et al. (*13*) note, even given the generous assumption that chromosomes 1A and 1B are harmless when present as a single copy, there will still be a strong negative frequency-dependent selection pressure against them, like any recessively lethal allele. Even if chance allowed the balanced lethal system to become fixed in a small sub- population, the ancestral version of chromosome 1 would be expected to rapidly invade upon secondary contact. In fact, the situation would appear still less favorable as, when combined with the ancestral chromosome, a single copy of chromosome 1A or 1B would be expected to be deleterious. Such a combination would possess only one copy of one of the deleted regions - exposing it to the effects of haploinsufficiency - while possessing three copies of the alternate region, leading to further unbalanced dosage. An analogue of this mixed genotype may be found in artificial crosses between *Triturus* and *Lissotriton*, which are occasionally viable to maturity, but with significantly reduced survival rates (*26*).

Counterintuitively, a resolution to this paradox can be found precisely *because* of the haploinsufficiency predicted for the mixed ancestral/rearranged genotype If an individual with the ancestral genotype disperses into a sub-population where, by chance, the balanced lethal system has become fixed, then all its offspring will carry one copy of the ancestral chromosome 1 and one copy of chromosome 1A or 1B (Fig. 4). These ‘hybrid’ offspring would suffer the unfavorable mixture of partially monosomy and trisomy described above. If the resultant fitness penalty is great enough, the offspring of the individual with the ancestral genotype will be less fit, on average, than offspring of parents that both carry the balanced lethal system. Therefore, within this sub- population, the ancestral un-rearranged form of chromosome 1 will be the one selected against most strongly. In this scenario, both the ancestral genotype and the balanced lethal system would experience positive frequency-dependent selection, with a tipping point above which the balanced lethal system will be driven towards fixation (Fig. 5A).

**Fig. 4:**
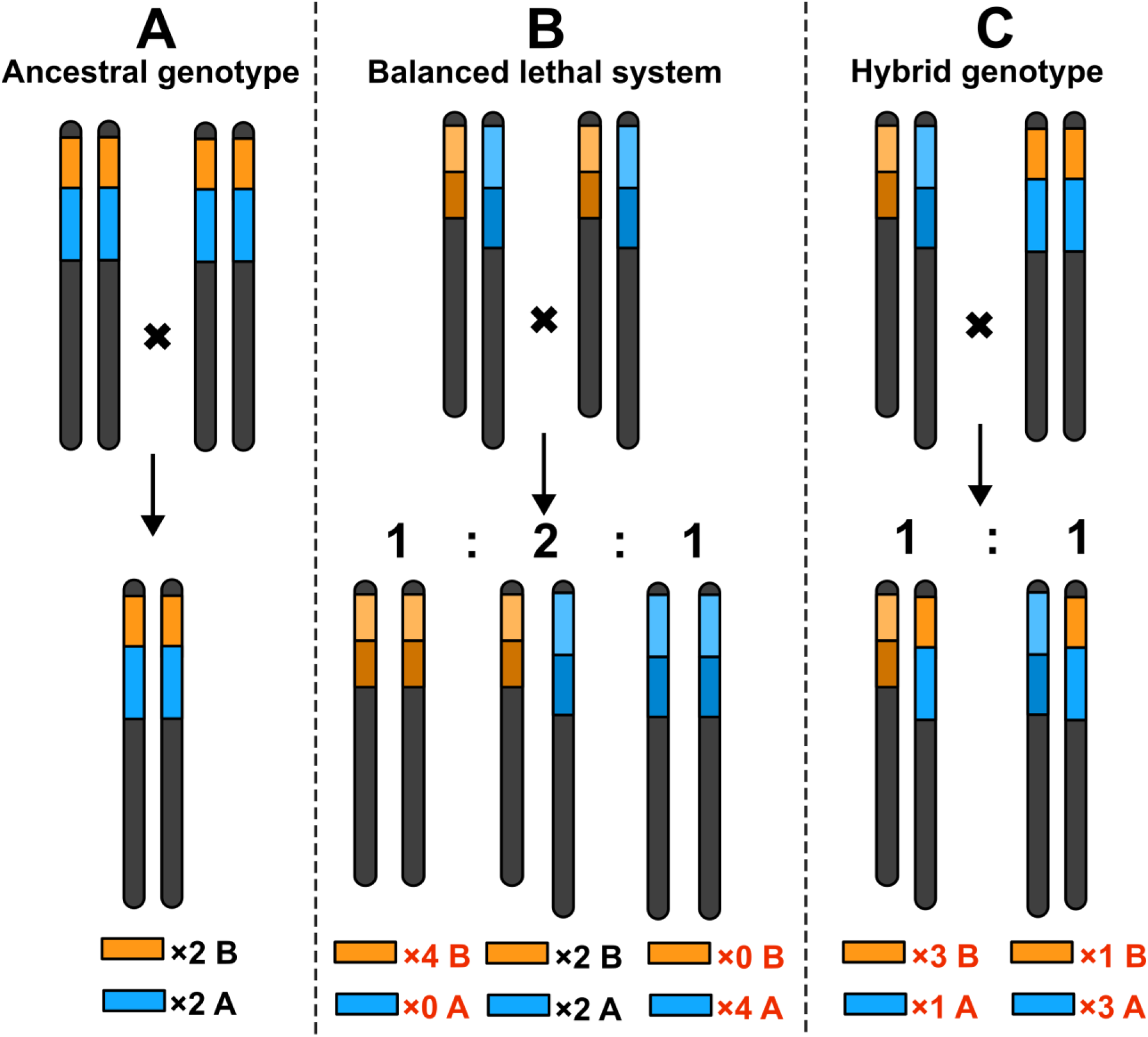
Consequences of the possible combinations of rearranged and ancestral genotypes. (**A**) In the case of a cross between two individuals of the ancestral (pre-rearrangement) genotype, all offspring will inherit two copies of all genes. (**B**) If both parents are affected by the balanced lethal system, then half of the offspring will inherit a full completement of genes, whereas the other half will have zero copies of either sets of genes involved and will thus not be viable. (**C**) In a cross between a parent affected by the balanced lethal system and a parent with the ancestral genotype, no offspring will inherit two copies of any of the genes involved in the balanced lethal system. Instead, these genes will be present in a mixture of single and triple copies, likely significantly reducing fitness.

**Fig. 5:**
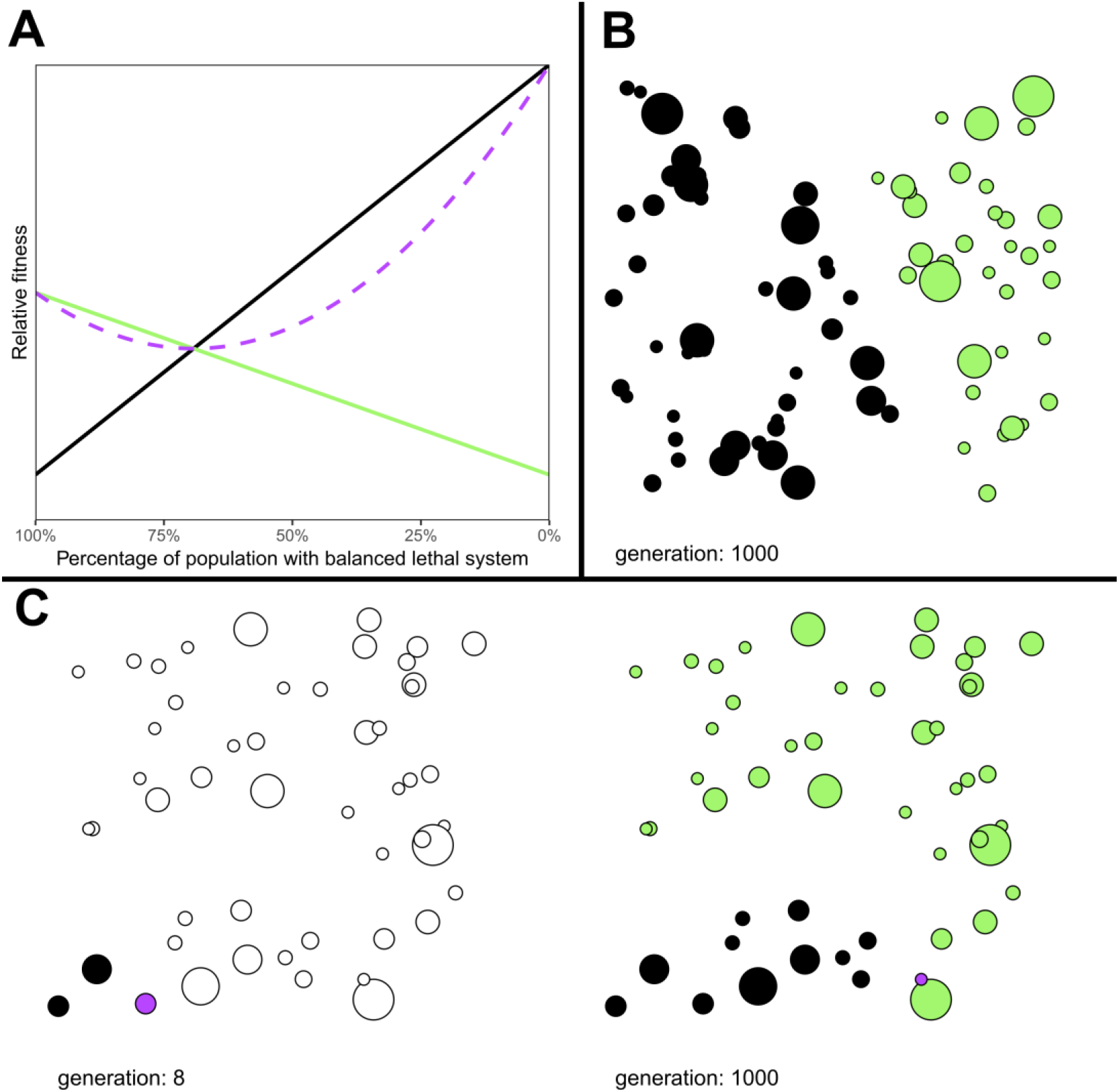
Models of outbreeding depression between the rearranged and ancestral genotypes. (**A**) Plot showing the relative fitness of offspring from parents with the ancestral pre-rearrangement (black) and balanced lethal (green) genotypes against the proportion of the ancestral chromosome in the population. If ‘mixed’ offspring (with one rearranged and one ancestral chromosome) have an average fitness lower than 0.5, there is a threshold below which the offspring of the parents of the balanced lethal system will have a higher fitness than those of the ancestral chromosome. The average fitness of offspring is shown by the dashed purple line with a minimum point, showing the effect of outbreeding depression. (**B**) Simulation of a persistent hybrid zone in a landscape of ponds (circles with area proportional to carrying capacity) formed after secondary contact between populations carrying the balanced lethal system (green) and ancestral chromosome (black), with ponds where neither genotype represents at least 80% of the population shown in purple, and empty ponds in white. Due to the lower fitness of the ‘mixed’ genotype, the ancestral chromosome is unable to displace the balanced lethal system (Fig. S6). (**C)** Simulation of the initiation of a balanced lethal system. An initial population (left) carrying the ancestral chromosome is founded at generation 0 in the pond in the bottom-left of the region. After 8 generations, when this population has grown to 317 individuals, a single migrant mutates to the balanced lethal genotype (resulting in the small purple pond). 1000 generations later (right), the region is split between the ancestral and balanced lethal genotypes, with a persistent hybrid zone formed (Fig. S7)

### Simulating the evolution of the balanced lethal system

We construct a model to explore whether heterozygote disadvantage between the derived and ancestral chromosomes stabilize a balanced lethal system. In this model the ancestral (NN) and heterokaryotic balanced lethal genotypes (AB) have equal fitness, the homokaryotic genotypes (AA and BB) are instantly lethal, and the mixed genotypes (AN and BN) have all fitness parameters reduced by 25% compared to the ancestral state. We simulate a scenario where two populations, one ancestral and one with the balanced lethal system fixed, colonize a region from opposite directions. In 42 of 100 replicates, we observe that after 1,000 generations of secondary contact a persistent hybrid zone has formed at a region of low population density (Figs. 5B, S6, Table S6). However, when this simulation is repeated with the mixed genotypes having full fitness, we observe that the balanced lethal system becomes extinct in all 100 replicates.

We then model the initiation of a balanced lethal system in a single individual. As Sessions et al. (*22*) note, if the rearrangement occurs early enough in an individual’s development, all gametes produced will carry either chromosome 1A or 1B. We simulate an expanding population with the ancestral genotype. After the adult population size has reached >200, a single individual on the periphery of this population has its genotype altered to AB. Over 50,000 replicates, we observe 109 (0.22%) instances of the formation of a persistent balanced lethal system after 1,000 generations (Figs. 5C S7, Table S6).

These simulations demonstrate that the unequal exchange, for which we find clear genomic evidence, is sufficient to explain both the origin and the persistence of the balanced lethal system in *Triturus*. As the initial mutation is present only in a single individual, the positive frequency dependent selection is strongly biased against it in the early generations. Therefore, the probability of the products of the unequal exchange surviving this period is very low – which accords with the fact that, aside from *Triturus*, there are no known examples of this genomic reconfiguration persisting. However, on the rare occasion that the products of the unequal exchange randomly become dominant in a small sub-population, then that sub-population can both persist and rapidly expand into uncolonized areas. This behavior may be characterized as genetic surfing (*27*), a phenomenon which can result in hybrid zones forming due to the effects of underdominant alleles during range expansion (*28*).

The structure of the *Triturus* genome matches the predicted outcome of an unequal exchange. However, we may consider whether any alternative mechanism could produce the same result. We observe four major mutations: two deletions and two duplications. Any scenario in which these each evolved independently is both less parsimonious than a single rearrangement and subject to mechanistic difficulties. Because any deletion of the size we observe in *Triturus* is almost certain to be lethal when homozygous, it could only be maintained at high frequency under powerful balancing selection. Effectively this requires that each deletion occurs within a pre-existing supergene system – for which we observe no evidence – and, even in this case, the deletion would still be disadvantaged in competition with the ancestral supergene. Evolution of a balanced lethal system in this scenario would require the sequential fixation of multiple, highly deleterious mutations within the entire population – without any equivalent of the reproductive isolation which protects the products resulting from an unequal exchange from being outcompeted.

## Conclusion

The balanced lethal system in *Triturus* newts is an anomaly of evolution in which an overwhelmingly deleterious trait persists, despite apparently offering no advantage. We reveal that the lethality and lack of recombination inherent in the system are caused by the total deletion of two large, adjacent sections of chromosome 1. We also show that both deletions are compensated for by a duplication of the same section on the opposite version of the chromosome. This is precisely the predicted outcome of an unequal exchange between chromosomes which, as Sessions et al. (*22*) presciently hypothesized, would form a balanced lethal system in a single step. As unequal recombination is often associated with repetitive elements (*29, 30*), it is perhaps no coincidence that this system has evolved in *Triturus* newts which, like other salamanders, have extremely large and repetitive genomes (*31*).

The duplications we observe allow chromosomes 1A and 1B to compensate for each other’s deficiencies, however ‘hybrids’, carrying one copy of the ancestral chromosome would still be affected by abnormal copy numbers of A- and B- linked genes. Due to this penalty, the balanced lethal system provides a relative fitness advantage in populations in which it is already dominant. When simulated, this mechanism allows the balanced lethal system to become fixed in a genetically distinct ‘population’, separated from its ancestor by a persistent ‘hybrid’ zone with severe outbreeding depression. This new ‘population’ is not only reproductively isolated, but also is characterized by a distinct phenotype (50% embryonic mortality) and thus subject to unique selective pressures – in effect, it behaves as a separate species. In this view, the balanced lethal system achieves fixation, not by spreading through an existing species, but by creating its own new species.

The architecture of the *Triturus* balanced lethal system provides an insight into how maladaptive traits can become fixed by reproductive isolation, through a newly described mechanism of instantaneous speciation via unequal exchange. It would be interesting to know whether this is a mechanism specific to *Triturus*, or if other naturally occurring balanced lethal systems (*6, 7*) have evolved in a similar way. Our work exemplifies that paradoxical evolutionary phenomena are worthy of special attention.

## Materials and Methods

All code used in this study can be found in the associated Zenodo repository (*32*), sequence data for all samples used is available at the Sequence Read Archive (see Table. S7 for accession numbers)

### Samples

For the *Triturus* linkage map a full-sibling family was bred at the University of Belgrade (Belgrade, Serbia). The experimental procedures were approved by the Ethics Committee of the Institute for Biological Research "Siniša Stanković”, University of Belgrade (decisions no. 03-03/16 and 01- 1949). The founder population consisted of two *T. macedonicus* males collected from Ceklin, Cetinje Municipality, Montenegro (42°21 N; 18°59 E) and two *T. ivanbureschi* females collect from Zli Dol, Pčinja District, Serbia (42°25 N; 22°27 E). Sampling from natural populations was approved by the Environmental Protection Agency of Montenegro (permit no. UPI-328/4) and Ministry of Energy, Development and Environmental Protection of the Republic of Serbia (permit no. 353-01-75/2014-08). From the F_0_ founders a male-female (non-sibling) pair of F_1_ *T. macedonicus* x *ivanbureschi* was raised to adulthood and mated producing 206 F_2_ *T. macedonicus* x *ivanbureschi* offspring (73 hatchlings and 133 arrested embryos). For the *Lissotriton* linkage map an analogous family was bred at Jagiellonian University (Kraków, Poland). Lissotriton samples were collected in accordance with the Polish General and Regional Inspectorates of Environmental Protection permits DOP-OZGIZ.6401.02.25.2011.JRO, OP-I.6401.32.2020.GZ, GDOŚ DZP-WG.6401.24.2021.TŁ and all experiments were accepted by the I Local Ethical Committee for Animal Experiments in Kraków, permit 28/2011 and the II Local Ethical Committee for Animal Experiments in Kraków, permit 64/2020. A non-sibling male-female pair of F_1_ *L. vulgaris* x *montandoni* was mated to produce 203 F_2_ offspring. Samples consisted of tale- tips taken from adult newts (the F_1_ parents of the *Triturus* and *Lissotriton* families, as well as the four F_0_ grandparents for *Triturus*), and whole embryos or hatchlings from offspring.

### DNA extraction, library preparation and target capture sequencing

Laboratory protocols followed the NewtCap protocol (*33*). Genomic DNA was extracted with the Promega Wizard^™^ Genomic DNA Purification Kit (Promega, Madison, WI, USA), according to the salt-based extraction protocol of Sambrook and Russel (*34*). 1,000 µg of DNA from each sample was used for library preparation, performed using the NEBNext Ultra^™^ II FS DNA Library Prep Kit for Illumina (New England Biolabs, MA, USA) following the protocol provided by the manufacturer, with all volumes divided by 4 and an enzymatic fragmentation time of 6:30 minutes. Size selection was performed using NucleoMag^™^ magnetic separation beads (Macherey-Nagel, Düren, Germany) targeting an insert size of 300 bp. Libraries were indexed via eight cycles of PCR, using unique combinations of i5 and i7 indices from IDT (Integrated DNA Technologies, Leuven, Belgium). Library concentration and fragment size distribution were measured via the Fragment Analyzer system (Agilent, Santa Clara, CA, USA) before the libraries were pooled equimolarly in batches of 16, with 250 ng of DNA per sample (4,000 ng total), and vacuum concentrated to 800 ng/µL.

Target enrichment was performed on the pooled libraries with the MyBaits-V4 kit (Arbor Biosciences, MI, USA). The bait set used (Ref# 170210-32) targets 7,139 genomic regions, based transcriptomes from multiple *Triturus* species (*35*). Enrichment followed the manufacturers protocol, with the following deviations: Blocks C and O were replaced with 30,000 ng of *Triturus* derived C_0_t-1 DNA to block the hybridization of repetitive sequences. Tissue to produce C_0_t-1 DNA was available from a removal action of an invasive population of *T. carnifex* (*36*). A hybridization time of 30 hours and temperature of 63 °C were employed, and libraries were incubated with the blocking solution for 30 minutes before addition of the RNA baits. After hybridization, the pools were amplified with 14 cycles of PCR before 150 bp paired-end sequencing, targeting a yield of 1 Gbp per sample, was performed on the NovaSeq 6000 platform (Illumina Inc., San Diego, CA, USA) by BaseClear B.V. (Leiden, the Netherlands).

### Processing of sequence capture data

Bioinformatics and analyses were performed in the Academic Leiden Interdisciplinary Cluster Environment (ALICE) at Leiden University. The upstream data processing was performed via a custom Perl (version v3.38.0) script (Pipeline_1.pl). FASTQ files containing demultiplexed raw sequence data were trimmed with Trimmomatic version 0.39 (*37*) and BBDuk version 38.96 (*38*) to remove adapter contamination. BWA-MEM version 0.7.17 (*39*) was used to map the trimmed reads against reference sequences previously assembled from *T. dobrogicus* (*35*). The resulting BAM files were processed, deduplicated and genotyped via the *AddOrReplaceReadGroups, MarkDuplicates* and *HaplotypeCaller* functions of GATK version 4.5.0.0 (*40*) producing a VCF file for each sample. Sequencing depth was assessed with a custom R (version 4.4) script (Peakloop2.R) which processed the output of the SAMtools version 1.18 (*41*) *depth* function of the deduplicated BAM file for each sample and evaluated the minimum depth of the best covered continuous 100 bp sequence within each target sequence of the reference assembly. All samples used for construction of the linkage maps were required to have a median best 100 bp sequencing depth of at least 10. After samples had been screened for coverage, each sample set was jointly genotyped with the *GenomicsDBImport* and *GenotypeGVCFs* function of GATK, producing a multi-sample GVCF file for each linkage family.

### *Lissotriton* linkage map construction

To construct the *Lissotriton* linkage map, VCFtools version 0.1.16 (*42*) was used to filter the multi-sample GVCF file to exclude indels and SNPs with a mean sequencing depth less than 10, genotype quality lower than 20, minor allele frequency lower than 0.4, or missing data greater than 5%. Finally, a single SNP per reference target was selected. A linkage map was then constructed with LepMAP3 version 0.5 (*43*). First the filtered GVCF was used as the input for the *ParentCall2* module, then initial linkage groups were created with the *SeparateChromosomes2* module, with a LOD limit of 22 and distortion LOD set to 1, unplaced markers were then incorporated with the *JoinSingles2All* module with a LOD limit of 15 (these settings were selected to maximize the numbers of included markers while yielding 12 linkage groups – the number of linkage groups increases rapidly as the LOD limit is raised until plateauing at 12 at LOD = 22, whereafter further increases only decrease the number of mapped markers). The markers were then ordered with the *OrderMarkers2* module, using 12 merge iterations and eight polish iterations, with the sexAveraged option enabled and the minError parameter set to 0.01. After construction the linkage groups were redesignated in order of decreasing length.

### Identification of chromosome 1 linked presence/absence markers in *Triturus*

The per-marker 100 bp peak region sequencing depth was used to identify presence/absence of markers in arrested embryos using a custom R script (Select_presence_absence_1.R). This first calculates expected sequencing depth scores for all markers across the sample set (based on the product of the mean depth per sample and the mean depth per marker), before identifying markers in which at least 25% of samples show a depth of zero, while an equivalent number of samples show more than double the expected depth (the predicated behaviour of presence/absence markers associated with either chromosome 1A or 1B). The scripts then clusters candidate presence/absence markers into sets of where the same samples show zero coverage. As at this stage the genotype of the arrested embryos was unknown, used the findings of de Visser, et al. (*21*) – based on phenotyped embryos - as a guide to designate each cluster as A- or B-linked, following the terminology of Macgregor & Horner (*4*) where 1A1A embryos are dubbed “slim-tailed” and 1B1B embryos “fat-tailed”.

### *Triturus* linkage map construction

The *Triturus* linkage map was constructed via the methodology described above, modified to incorporate the presence/absence markers. To this end the coverage scores of these markers were converted into probabilistic pseudo-SNP calls using a custom R script (Add_presence_absence_to_call_table_4.R). For example, where the measured coverage was close to expected, the genotype could be expressed as GT, where there was zero coverage GG, and where there was twice the expected coverage TT. These calls where then appended to the output of the *ParentCall2* module of LepMAP3 (*43*). From this point the LepMAP3 pipeline was used with the same settings as with *Lissotriton*, with the expected that the LOD limits for the *SeparateChromosomes2* and *JoinSingles2All* modules were changed to 27 and 20 respectively (as with *Lissotriton* these settings were chosen as the point where the number of linkage groups plateaus at 12). After construction of an initial map, the linkage group where the presence/absence markers clustered was assigned linkage group 1 and additional, separate maps for chromosome 1A and 1B (each including only one of the two sets of presence/absence markers) were constructed. The remaining linkage groups were ordered by length.

### Comparison with *P. waltl* and *A. mexicanum* genome assemblies

The *Triturus* reference sequences placed on the linkage map were aligned against the against the genome assemblies published for the Iberian ribbed newt, *Pleurodeles waltl* (*17*) and the axolotl, *Ambystoma mexicanum* (*18, 19*) using BLAST+ version v2.14.1 (*44*). For *P. waltl* the default setting were used except for specifying a minimum E-value of 1e-20 for the more distantly related *A. mexicanum* a minimum E-value of 1e-10 and word size of 15 was used. Sequences that aligned outside of the main chromosomes of the assemblies or aligned in multiple locations were removed.

### Ploidy analysis

Data from thirty F_1_ *T. macedonicus* x *T. ivanbureschi* samples from a previous study (*25*) (10 each of genotype AA, AB and BB) were used to examine SNP allele ratios of the A and B-linked genes. F_1_ hybrids are desirable for this application as their use maximizes the number of heterozygous SNPs. These samples were processed with the same target capture methodology as the linkage map samples. The BAM files produced for each sample were subsetted to produce one file with alignments for all 7,139 references sequences, and two each with alignment for 28 A-linked genes and 35 B-linked genes - selected as targets in which presence/absence variation has been validated across the genus (*21*). These alignments were processed with nQuire (cloned from Git commit 8975c94) (*45*), using the *create* function with -c set to 20 and -p to 10. The *denoise* function was then used on the resulting .bin files followed by ploidy model fitting with the *histotest* and *view* functions. The resulting allele ratio distributions were normalized and combined. The *R*^*2*^ values comparing the fit of the observations to calculated allele distributions were averaged for each sample genotype and gene-category. Significant of agreement of best fit model was calculated with p ≤ 0.05 requiring at least eight out the ten individuals in each genotype category to agree. For additional context mean coverage data from the same gene and genotypes categories of the F_2_ *T. macedonicus* x *ivanbureschi* sample set calculated.

### Simulation of balanced lethal system evolution

A custom model, BL_sim_3.R, was developed in R. To examine whether the balanced lethal system could be protected from invasion by the ancestral chromosome by the effects of heterozygote disadvantage 100 replicates of this simulation were run for 1,000 generations in secondary contact mode with the hybrid fitness parameter set to its default value of 0.75. This simulation was then repeated with hybrid fitness set to 1.0 (simulating the effects of no outbreeding depression). To examine the fate of an unequal exchange we then simulated 50,000 replicates (spread across 5 runs) of the model in its single mutation mode, with hybrid fitness parameter set to 0.75 and a max run time of 1,000 generations.

## Acknowledgments

We thank R. Butlin, W. Halfwerk and M. Yun for their insightful discussion during this study. We thank M. Fahrbach for the images used in Fig. 1 and E. Bosch for the illustrations used in Fig. 2 (licensed by Naturalis - CC-BY-NC-ND 4.0). We thank the staff of Leiden University’s ALICE cluster for their technical support

## Author contributions

Conceptualization: PA, WB, AI, JS, BW

Breeding of Triturus linkage family: MC, AI, TV

Breeding of Lissotriton linkage family: WB

Target capture sequencing: JF

Linkage map analysis: JF

Ploidy analysis: MCdV

Evolutionary modelling: JF

Manuscript preparation: JF, BW with assistance from all other authors

## Data and materials availability

All raw sequence data used in this study is available via at the Sequence Read Archive (see Table S7 for accession numbers). All code used in this study as well as markdown documents detailing all commands used in the workflow are archived at this studies Zenodo repository (*32*), together with an .xlsx document detailing the sequences and positions of all markers located on the *Triturus* and *Lissotriton* linkage maps.

## Supplementary Materials

Supplementary Text – Description of Simulation

Figs. S1 to S7

Tables S1 to S7

