## Supplementary Data for "The balanced lethal system in *Triturus* newts originated in an instantaneous speciation event"

**Description of BL_sim_3.R**

BL_sim_3.R is a non-Fisher-Wright model, featuring overlapping generations and local colonization and extinction. In each run of the script the requested number of replicates will be simulated under the requested conditions. The outcome of each run is determined by the conditions, and a random seed (may be input by the user in order to repeat a run with the exact same outcome).

For each replicate of the simulation, a landscape containing 50 to 150 breeding localities (i.e., ponds) is randomly generated in a 5×5 km area, with each pond given a size score (determining carrying capacity) between 5 and 60, following a power law distribution. The simulation cycles through three phases. In the breeding phase, each adult female (of minimally two years of age) chooses a male in the same pond as her mate randomly, weighed according to male attractiveness and produces a number of embryos equal to her fecundity. If the adult population exceeds the ponds size value each adult is given an increasing chance to skip the breeding phase, such that on average the breeding population is limited to the size of the pond – selectively skipping breeding seasons is common in *Triturus* newts (*46*) and other amphibians (*47*). The embryos' genotype is a random combination of their parents and determines the chance of the embryo hatching. After hatching a randomly selected 95% of individuals are culled to simulate larval mortality. In the dispersal phase all newts of age 1 have a 50% chance of moving to another pond within 1 km, the destinations are selected randomly and weighted linearly with increasing size and decreasing distance. In the aging phase each individual has a survival chance determined by age and genotype, and survivor age is incremented by 1.

All individuals have a genotype with three available alleles, N represents the ancestral chromosome where A and B are the heteromorphs of the balanced lethal system. Genotypes NN and AB have values for embryonic survival, juvenile survival, adult survival, female fecundity and male attractiveness of 1.0, 0.2, 0.8, 200 and 1.0 respectively. The survival and fecundity values are based on studies on *Triturus cristatus* by Arntzen and Teunis (*48*) and modelling by Griffiths and Williams (*49*). As it is unknown at which life stage(s) a hybrid fitness penalty will be most significant, it is modelled as a proportional reduction across all these parameters. Male attractiveness is incorporated into balance the fitness penalty to fecundity – which would otherwise result in a bias where female hybrids have lower relative fitness than males. Genotypes BB and AA have values of zero for all parameters, whereas hybrid ancestral/balanced lethal system genotypes (AN and BN) are given values of X × that of genotype NN, where X is an input given by the user. The default value of X is (arbitrarily chosen as) 0.75, so the resulting hybrid fitness parameters are (0.75, 0.15, 0.6, 150 and 0.75).

Note that in this model fitness (expected number of offspring from each individual) is proportional to each of the embryonic survival, juvenile survival, and fecundity (or attractiveness in males) parameters. Therefore, a $\frac{1}{4}$ reduction across all three will result in a relative fitness of $(\frac{3}{4})^3 = \frac{27}{64}$. Adult mortality is reapplied every cycle of the simulation. Consequently, fitness is proportional to the sum of the geometric series $S^0 + S^1 + S^2 + \cdots + S^n = \frac{1}{1-S}$ where $S$ is the adult survival parameter. Therefore, with the adult survival parameter for ancestral (NN) and heterozygote balanced lethal (AB) genotypes set as 0.8 and adult survival set to 0.6 for (AN and BN) hybrids the relative fitness of hybrids is multiplied by $\frac{1}{1-0.6} / \frac{1}{1-0.8} = \frac{1}{2}$ for a final relative fitness of $\frac{1}{2} \times \frac{27}{64} = \frac{27}{128} \approx 0.211$

The simulation allows for two basic scenarios. In secondary contact mode, two populations (each starting with of 20 adult individuals) one with the balanced lethal system fixed, the other carrying the ancestral chromosome, are initialized at opposite ends of the landscape and allowed to spread towards each other. In single mutation mode a single population with the ancestral chromosome fixed is initialized at the edge of the landscape and allowed to reproduce until reaching a population of at least 200 adults, at which point a single individual is randomly chosen to have its genotype flipped to AB. After each run of the simulation is completed the results (the locations of, and population history of each pond) are saved and then may be visualized with the accompanying script Visulise_2.R.

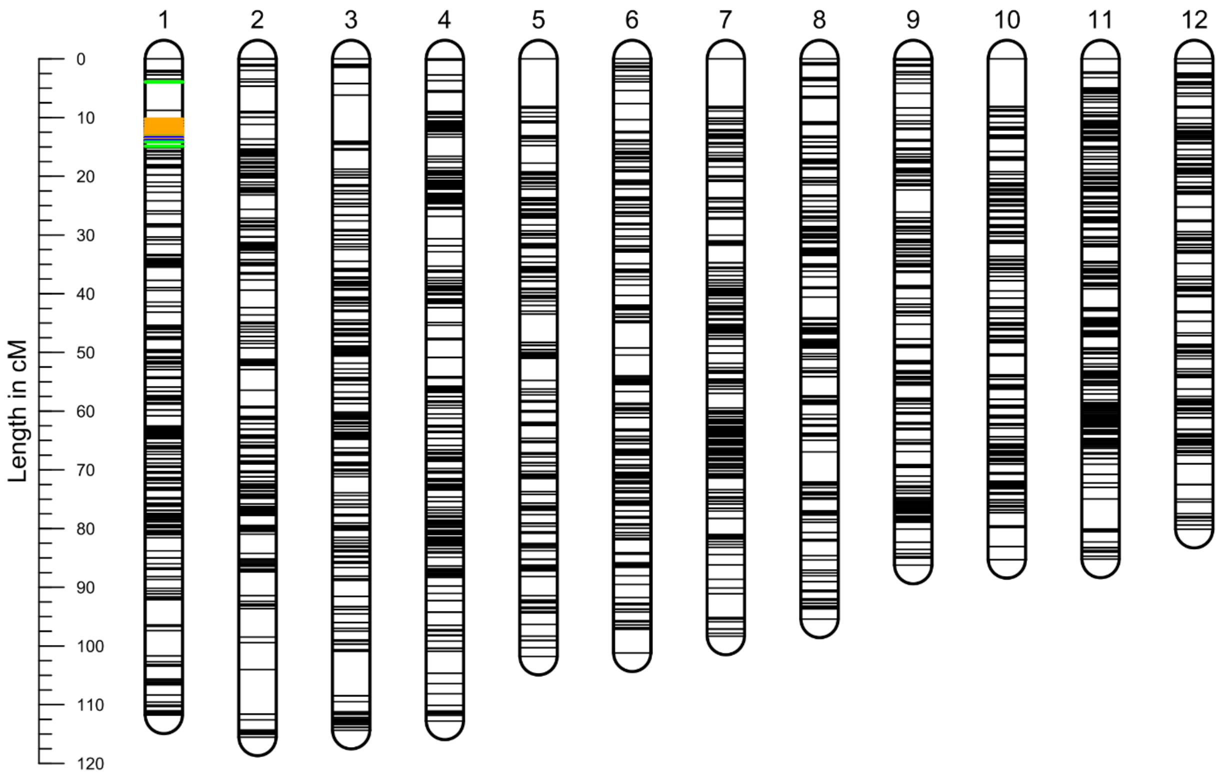


**Fig. S1. *Triturus* Linkage map.** Based on target capture data from a full-sibling $F_2$ *T. ivanbureschi* × *macedonicus* family, consisting of four $F_0$ grandparents, two $F_1$ parents and 206 $F_2$ offspring. The map includes 4,226 markers in 12 linkage groups, spanning a total length of 1,188 cM. Linkage groups are arranged by length, except for group 1, which is designated in accordance with the chromosome 1 linked presence/absence markers we located within it. Group 1 includes 29 A-linked markers, highlighted in blue and 33 B-linked makers highlighted in orange, 43 of these markers map to a single position at 13.659 cM from the group's origin, with the reminder deviating by up to 2 cM (for a more detailed view of group 1, split into chromosome 1A and 1B, see Fig. *2B*) .This deviation is likely an artifact caused by translating the presence/absence data of low coverage markers into pseudo-SNP calls: with perfect data we would expect all markers to collapse to a single point. An additional 12 markers, highlighted in green, were found associated with the balanced lethal systems in other *Triturus* species in a separate study (*21*) – ten of these cluster within 1 cM of the presence/absence markers, while the other two are displaced by approximately 10 cM, possibly because they show presence/absence variation in *T. macedonicus* but not *T. ivanbureschi*.

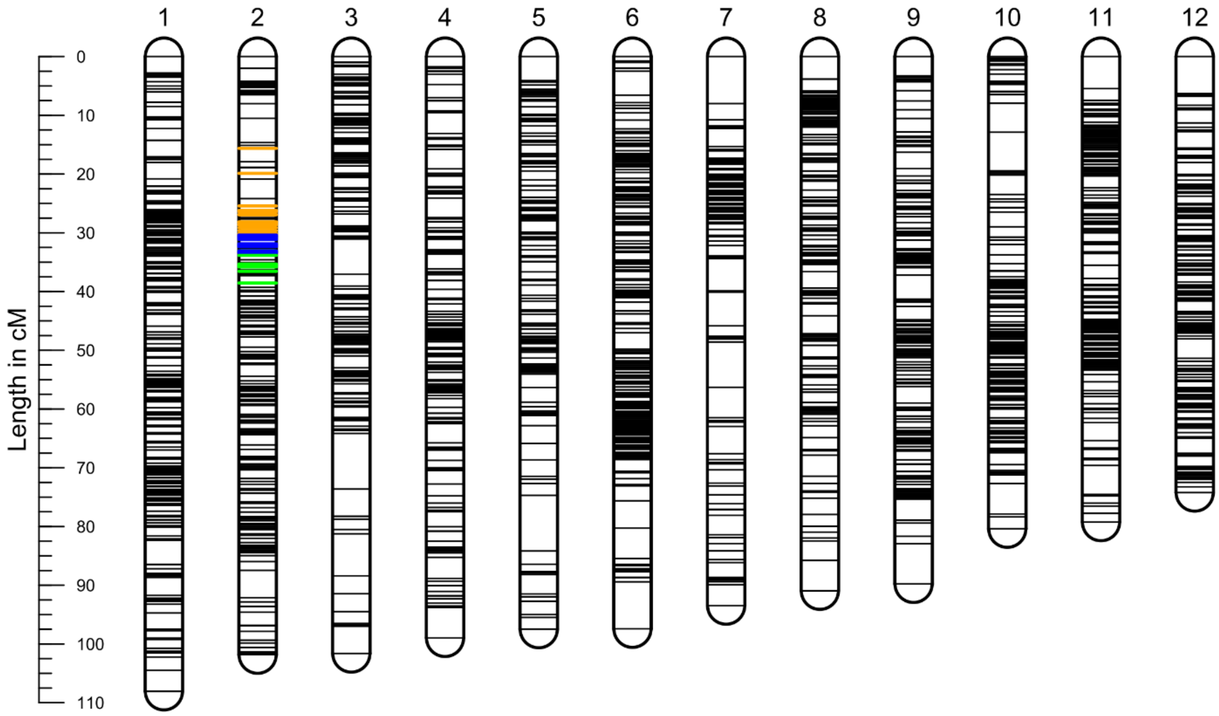


**Fig. S2. *Lissotriton* linkage map**. Based on target capture data from a full-sibling F2 *Lissotriton vulgaris* × *montandoni* family, consisting of two F1 parents and 203 F2 offspring. The map includes 3,693 markers in 12 linkage groups, spanning a total length of 1,113 cM. In *Lissotriton* the homologs of genes associated with the *Triturus* balanced lethal system are found in linkage group 2, where they form distinct blocks, corresponding to genes present only in *Triturus* chromosome 1A or 1B (highlighted in blue and orange). A third block includes genes (highlighted in green) that show either species-specific presence/absence variation within *Triturus* (*21*), or extreme heterozygosity specifically in viable embryos - indicating there are two distinct alleles, each associated with only one form of *Triturus* chromosome 1.

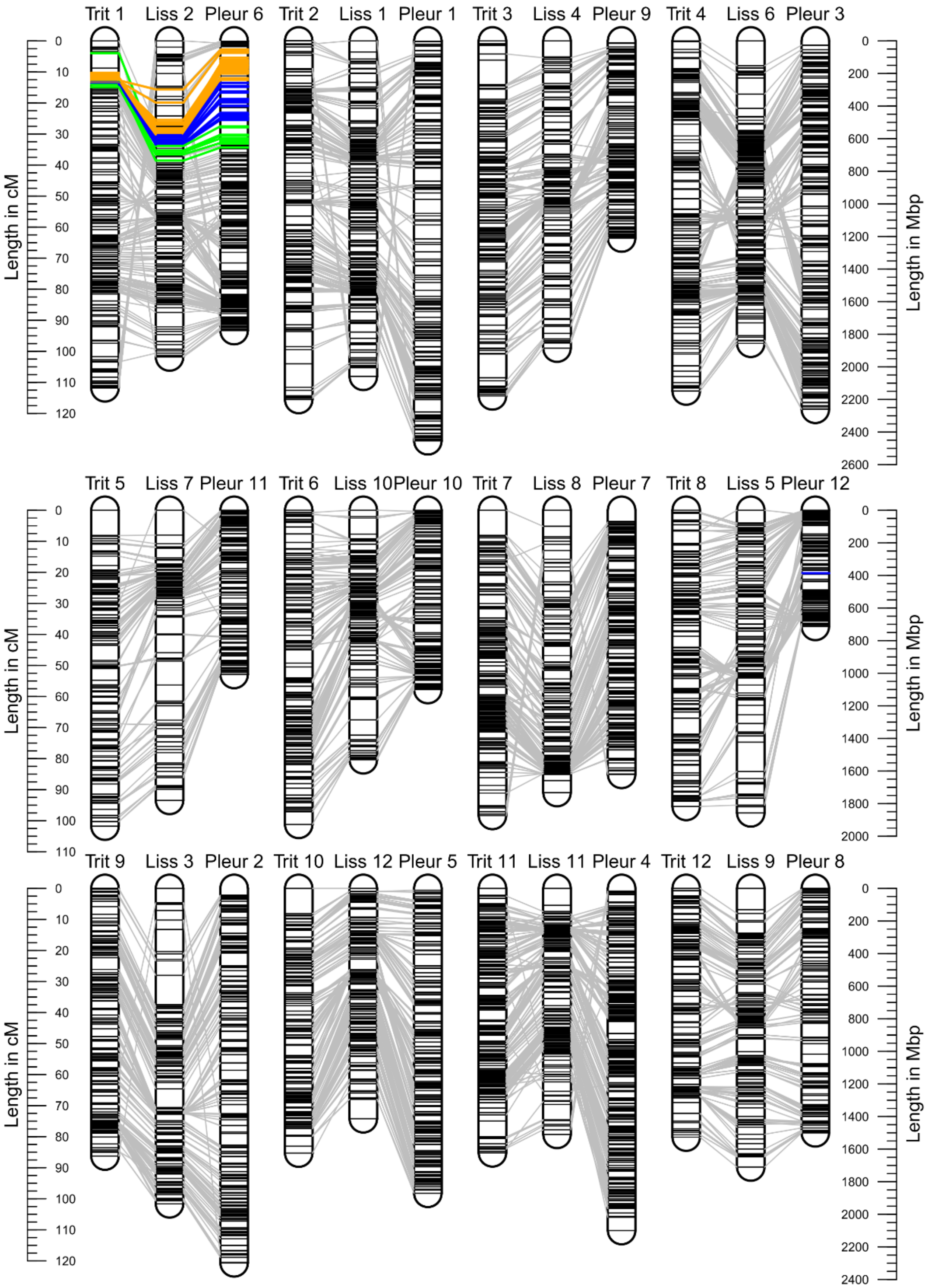


**Fig. S3. Homology between three newt genera.** The *Triturus* (Trit) and *Lissotriton* (Liss) linkage maps and the *Pleurodeles* (Pleur) genome assembly (*17*) show tight conservation of synteny. *Triturus* chromosome 1 linked markers are highlighted: A-linked in blue, B-linked in orange and species-specific in green.

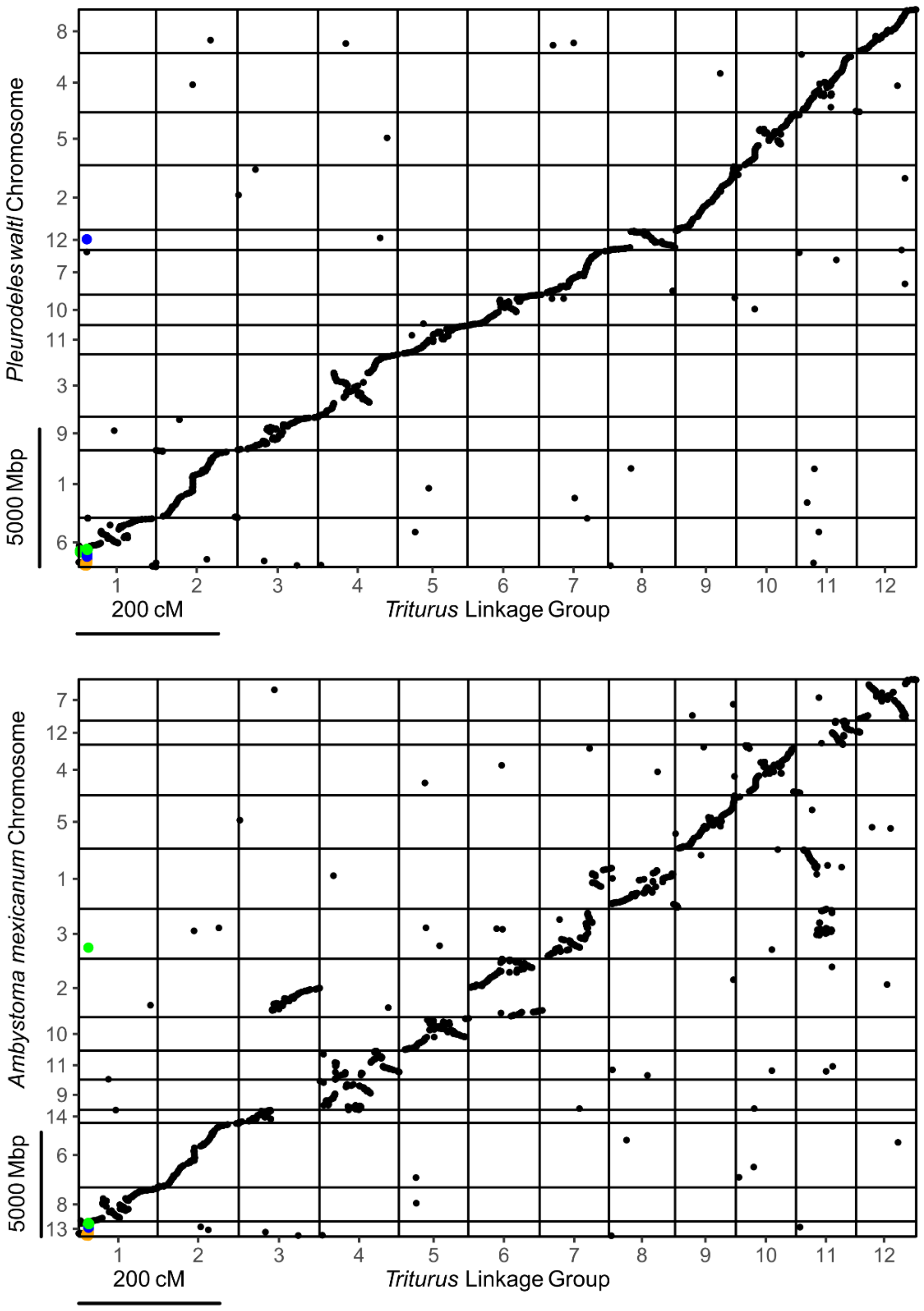


**Fig. S4. Oxford plots of the *Triturus* linkage map.** Showing locations of markers within the linkage map and genome assemblies for *Pleurodeles* (*17*) and *Ambystoma* (*18*, *19*). Synteny is extremely tightly conserved between the two newt taxa, but less so in the more distantly related *Ambystoma*.

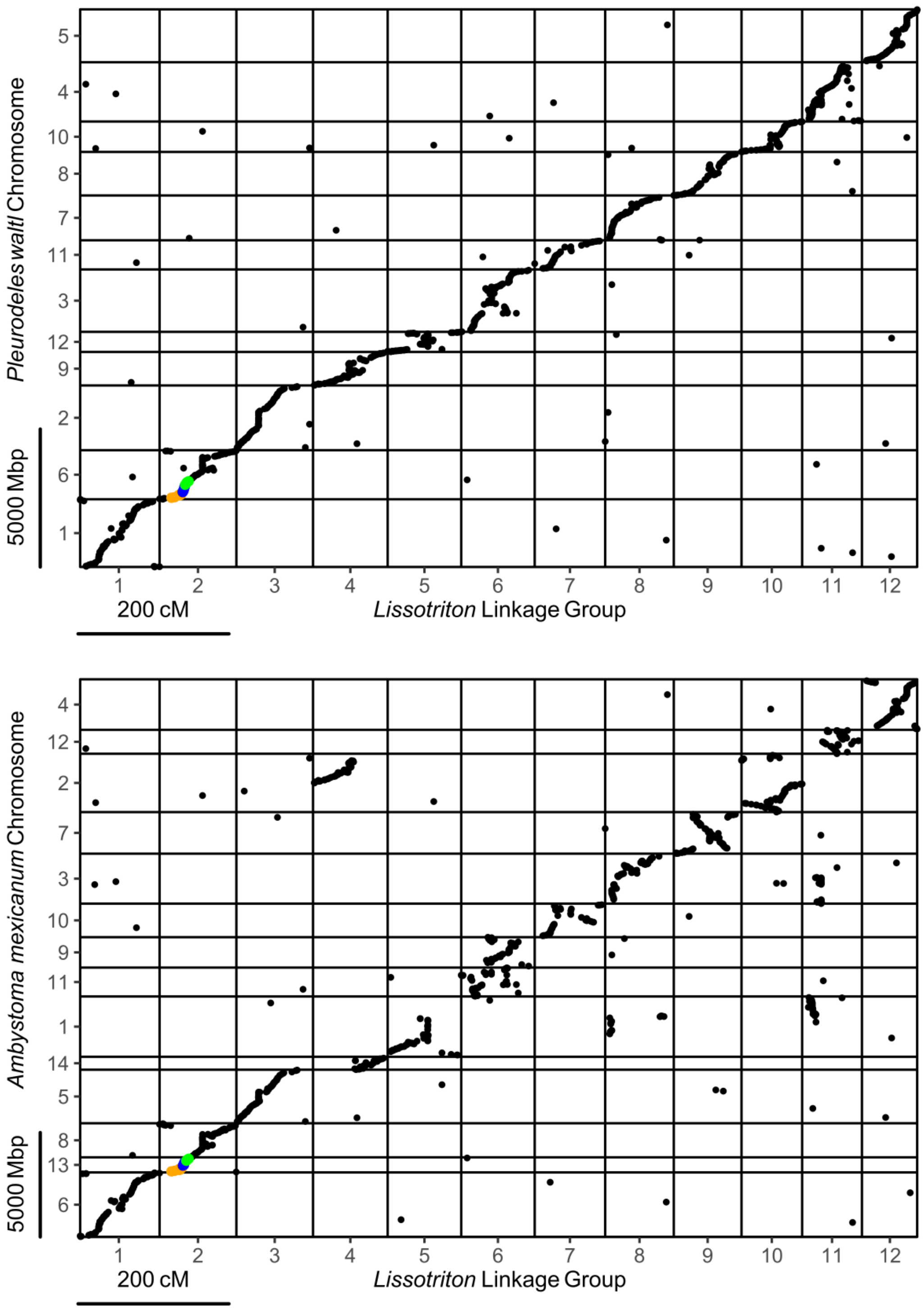


**Fig. S5. Oxford plots of the *Lissotriton* linkage map**. Syntenny is also tightly conserved between the *Lissotriton* linkage map and genome assemblies for *Pleurodeles* and *Ambystoma* (*17*–*19*).

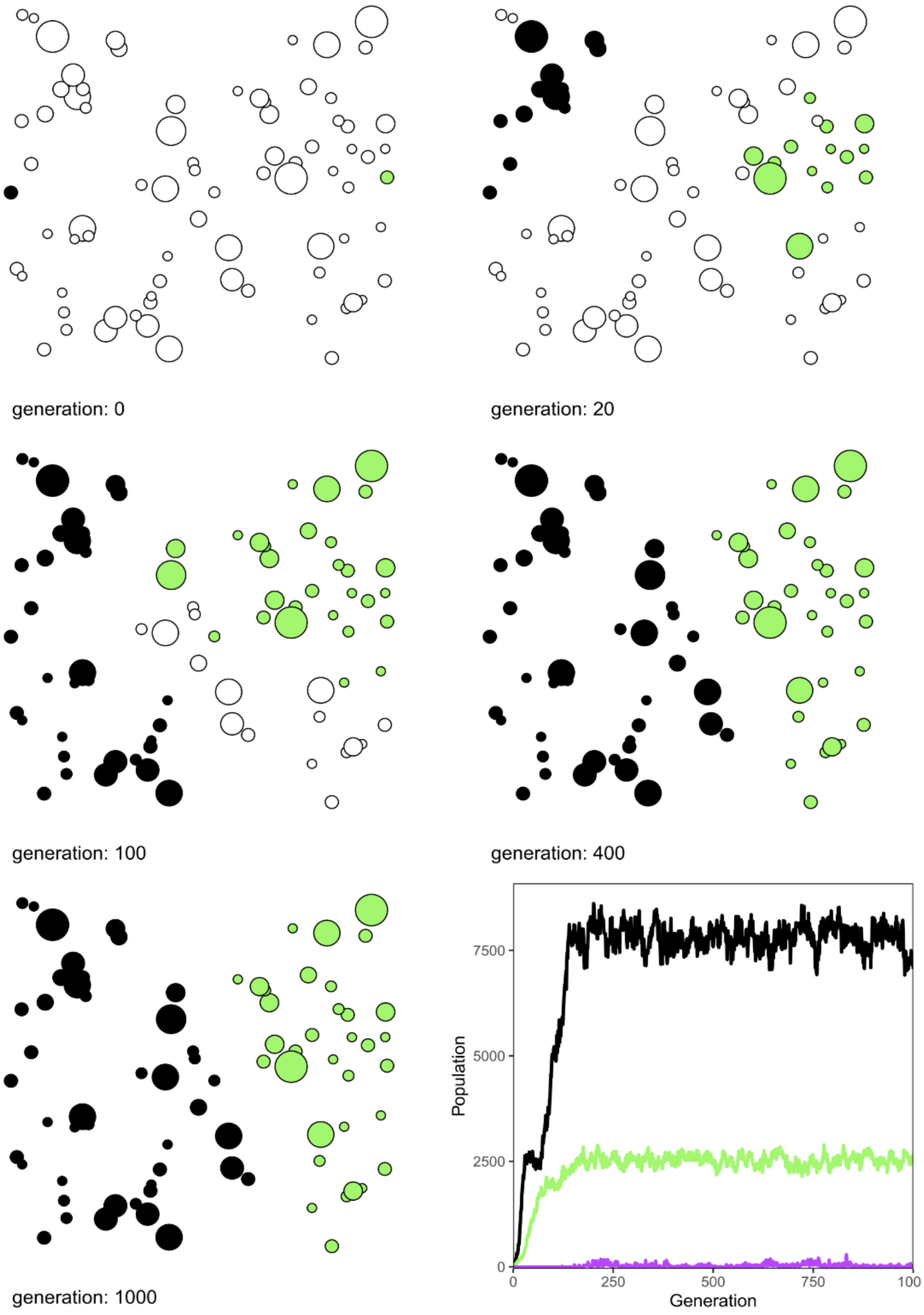


**Fig. S6. Simulation of secondary contact.** A population with the ancestral genotype (black) initiated at the left edge of the area expands towards a (green) population with the balance lethal system fixed. Despite the apparent fitness disadvantage of the balanced lethal population the effects of underdominance in chromosome 1 allows a persistent hybrid zone to form in an area that has low potential population density. Populations plateau after all ponds are colonized at approximately generation 200. Mixed populations and hybrid individuals are indicated in purple. (this example is the 19$^{th}$ replicate generated with BL_sim_3.R in secondary contact mode with hybrid fitness parameters set to 0.75 starting from a random seed of 6122263)

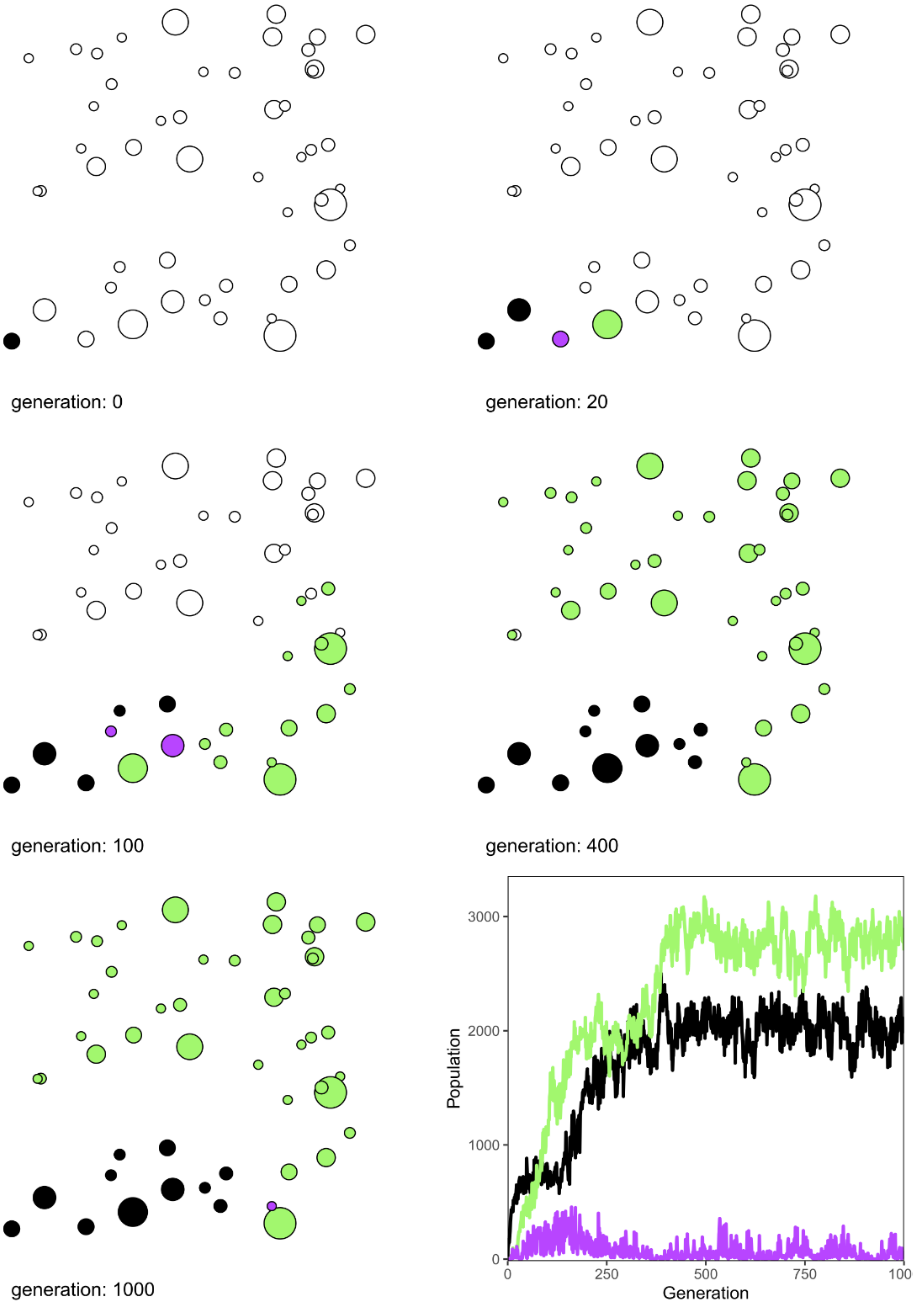


**Fig. S7. Origin of a balanced lethal system.** A population with the ancestral genotype (black) is initiated in the bottom left corner of the area. At generation 8, when the population size is 317 a single individual has its genotype altered to AB. Random chance allows the balanced lethal system (indicated in green, with mixed populations and hybrids indicated in purple) to colonize in the direction of a geographic bottleneck, which becomes the location of a persistent hybrid zone separating the two populations. (this example is the 978th replicate generated with BL_sim_3.R in single mutation mode with hybrid fitness parameters set to 0.75 starting from a random seed of 1088689)

| | *Lissotriton* | | *Triturus* | |
|---|---|---|---|---|
| **Group** | **Number of markers** | **Length (cM)** | **Number of markers** | **Length (cM)** |
| **1** | 410 | 108.1 | 399 | 111.8 |
| **2** | 343 | 101.8 | 335 | 115.5 |
| **3** | 257 | 101.6 | 339 | 114.4 |
| **4** | 259 | 99.0 | 471 | 112.8 |
| **5** | 239 | 97.5 | 290 | 101.8 |
| **6** | 465 | 97.4 | 299 | 101.2 |
| **7** | 167 | 93.5 | 388 | 98.3 |
| **8** | 302 | 90.9 | 284 | 95.4 |
| **9** | 263 | 89.8 | 322 | 86.2 |
| **10** | 305 | 80.4 | 309 | 85.3 |
| **11** | 409 | 79.3 | 518 | 85.2 |
| **12** | 274 | 74.2 | 272 | 80.1 |
| **Total** | **3693** | **1113.5** | **4226** | **1188.1** |

**Table S1. Characteristics of the linkage maps.** The target capture linkage maps produced for *Triturus* and *Lissotriton* both consist of 12 linkage groups, matching the karyotypes of both genera.

| | *Triturus* Linkage map | |
|---|---|---|
| | **Common Loci** | **Loci on Homologous Chromosomes** |
| ***Lissotriton*** | 2551 | 2497 (97.8%) |
| ***P. waltl*** | 3400 | 3359 (98.7%) |
| ***A. mexicanum*** | 2816 | 2408 (85.5%) |

**Table S2. Common loci of the *Triturus* linkage map.** The number of loci identified on the *Triturus* linkage map compared to the number of homologs found within the *Lissotriton* linkage map and the *Pleurodeles waltl* (*17*) and *Ambystoma mexicanum* (*18, 19*) genome assemblies (out of a total of 4226 loci placed on the *Triturus* map). The proportion of loci mapping to homologous chromosomes is very high in *Lissotriton* and *P. Waltl*, but lower in *A. mexicanum* due to several rearrangements.

| | ***R^2* scores** | | | | | | | | |
|---|---|---|---|---|---|---|---|---|---|
| **Sample and Genotype** | **All Markers (n = 7139)** | | | **A-linked Markers (n = 28)** | | | **B-linked Markers (n = 35)** | | |
| | **Diploid** | **Triploid** | **Tetraploid** | **Diploid** | **Triploid** | **Tetraploid** | **Diploid** | **Triploid** | **Tetraploid** |
| BW_0024 **AB** | **0.94** | *0.36* | *0.01* | **0.26** | *0.01* | *0.20* | **0.48** | *0.03* | *0.05* |
| BW_0025 **AB** | **0.91** | *0.32* | *0.00* | **0.61** | *0.10* | *0.03* | **0.16** | *0.00* | *0.20* |
| BW_0026 **AB** | **0.93** | *0.36* | *0.01* | **0.40** | *0.19* | *0.06* | **0.27** | *0.00* | *0.08* |
| BW_0027 **AB** | **0.94** | *0.30* | *0.00* | **0.50** | *0.16* | *0.07* | **0.28** | *0.00* | *0.15* |
| BW_0028 **AB** | **0.88** | *0.29* | *0.00* | **0.26** | *0.03* | *0.13* | **0.23** | *0.02* | *0.08* |
| BW_0029 **AB** | **0.85** | *0.31* | *0.00* | *0.00* | *0.00* | **0.09** | **0.43** | *0.19* | *0.01* |
| BW_0030 **AB** | **0.78** | *0.20* | *0.01* | **0.35** | *0.17* | *0.02* | **0.46** | *0.09* | *0.02* |
| BW_0031 **AB** | **0.83** | *0.28* | *0.00* | **0.27** | *0.23* | *0.00* | **0.43** | *0.05* | *0.09* |
| BW_0051 **AB** | **0.92** | *0.34* | *0.00* | **0.61** | *0.07* | *0.01* | **0.38** | *0.02* | *0.01* |
| BW_0052 **AB** | **0.91** | *0.32* | *0.00* | *0.09* | *0.00* | **0.10** | **0.52** | *0.02* | *0.11* |
| BW_0040 **AA** | **0.45** | *0.13* | *0.01* | *0.04* | *0.02* | **0.07** | | NA | |
| BW_0041 **AA** | **0.38** | *0.06* | *0.02* | *0.11* | **0.38** | *0.08* | | NA | |
| BW_0042 **AA** | **0.46** | *0.16* | *0.01* | **0.03** | *0.01* | *0.01* | | NA | |
| BW_0043 **AA** | **0.72** | *0.20* | *0.01* | **0.07** | *0.05* | *0.04* | | NA | |
| BW_0044 **AA** | **0.94** | *0.37* | *0.01* | *0.02* | *0.13* | **0.22** | *0.03** | *0.00** | *0.01** |
| BW_0045 **AA** | **0.94** | *0.31* | *0.01* | **0.21** | *0.17* | *0.00* | | NA | |
| BW_0046 **AA** | **0.91** | *0.32* | *0.00* | *0.01* | **0.02** | *0.01* | | NA | |
| BW_0047 **AA** | **0.90** | *0.34* | *0.00* | **0.02** | *0.01* | *0.00* | | NA | |
| BW_0064 **AA** | **0.90** | *0.35* | *0.00* | *0.04* | *0.15* | **0.22** | *0.02** | *0.03** | *0.09** |
| BW_0065 **AA** | **0.93** | *0.34* | *0.01* | *0.08* | *0.03* | **0.24** | | NA | |
| BW_0032 **BB** | **0.53** | *0.22* | *0.01* | | NA | | **0.16** | *0.00* | *0.01* |
| BW_0033 **BB** | **0.47** | *0.10* | *0.05* | | NA | | *0.02* | *0.01* | **0.19** |
| BW_0034 **BB** | **0.57** | *0.18* | *0.00* | | NA | | *0.02* | *0.02* | **0.03** |
| BW_0035 **BB** | **0.42** | *0.11* | *0.03* | | NA | | **0.10** | *0.05* | *0.01* |
| BW_0036 **BB** | **0.83** | *0.31* | *0.00* | | NA | | **0.38** | *0.01* | *0.00* |
| BW_0037 **BB** | **0.93** | *0.42* | *0.03* | | NA | | *0.20* | *0.19* | **0.21** |
| BW_0038 **BB** | **0.91** | *0.33* | *0.00* | | NA | | *0.02* | **0.25** | *0.08* |
| BW_0039 **BB** | **0.93** | *0.36* | *0.01* | | NA | | *0.01* | **0.20** | *0.11* |
| BW_0056 **BB** | **0.92** | *0.35* | *0.00* | | NA | | *0.00* | *0.02* | **0.04** |
| BW_0057 **BB** | **0.95** | *0.37* | *0.02* | | NA | | *0.01* | **0.07** | *0.03* |

**Table S3. Per sample results of ploidy analysis.** $R^2$ values are calculated showing the best fit nQuire (*45*) ploidy model (highlighted in green with bold text) for all genes, A-linked genes and B-linked genes in 30 $F_1$ *Triturus ivanbureschi* × *macedonicus* samples (*25*), split evenly between the three chromosome 1 genotypes. For BB samples no A-linked loci whatsoever were available for analysis. In two AA samples $R^2$ values (marked with *) could be calculated for reads mapped to a single B-linked marker, but this is not sufficient data to produce a meaningful result, and these reads likely represent artifacts.

| Sample Genotype | All Markers | | A-linked Markers | | B-linked Markers | |
|---|---|---|---|---|---|---|
| | Total Genes | Total SNPs | Total Genes | Total SNPs | Total Genes | Total SNPs |
| **AB** | 4539.2 | 12484.5 | 15.9 | 166.6 | 19.1 | 168.4 |
| **AA** | 4130.4 | 10756.1 | 22.8 | 123.2 | 0.2 | 0.5 |
| **BB** | 4055.8 | 10574.1 | 0.0 | 0.0 | 22.4 | 104.0 |

**Table S4. Details of SNPs used to analyze ploidy.** Average number of markers and SNPs (averaged across the 10 $F_1$ *Triturus ivanbureschi* × *macedonicus* samples (*25*) of each genotype) used to calculate $R^2$ values for ploidy models for each of three categories of markers.

| Sample Genotype | Coverage: All Markers | | Coverage: A-linked Markers | | Coverage: B-linked Markers | |
|---|---|---|---|---|---|---|
| | Raw | Normalized | Raw | Normalized | Raw | Normalized |
| **AB** | 58.6 | 58.6 | 42.6 | 42.6 | 32.1 | 32.1 |
| **AA** | 74.4 | 58.6 | 107.3 | 84.5 | 0.1 | 0.1 |
| **BB** | 45.2 | 58.6 | 0.1 | 0.1 | 53.8 | 69.7 |

**Table S5. Details of coverage in linkage map data set.** Mean coverage (measured at the best covered 100 bp sequence within each marker) of the 206 $F_2$ *Triturus ivanbureschi* × *macedonicus* offspring of the linkage map family, aggregated by marker category and sample genotype. The ratio of mean raw coverage across all markers was used to normalize the mean coverage of the three genotypes. Coverage of A-linked markers in samples of genotype AA is approximately double that of these markers in samples of genotype AB, and the same applies to B-linked markers in samples of genotype BB.

| Scenario | Hybrid Fitness | Random Seed | Replicates | AB survival |
|---|---|---|---|---|
| Secondary contact | 0.75 | 6122263 | 100 | 42 |
| Secondary contact | 1 | 4037782 | 100 | 0 |
| Single mutation | 0.75 | 1088689 | 10000 | 23 |
| Single mutation | 0.75 | 6036874 | 10000 | 23 |
| Single mutation | 0.75 | 4955070 | 10000 | 25 |
| Single mutation | 0.75 | 96996 | 10000 | 17 |
| Single mutation | 0.75 | 8259011 | 10000 | 21 |

**Table S6**. **Simulations of balanced lethal system.** Simulation of secondary contact was performed in two runs of 100 replicates. In the first run, with hybrid fitness set to its default value of 0.75 the balanced lethal system survived 1,000 generations of secondary contact in 42 replicates (e.g. Fig. S6). In the second run hybrid fitness was set to 1.0, and without the outbreeding depression to protect it the balance lethal system was extinct by generation 1,000 in all 100 runs. Simulation of the unequal exchange giving rise to the balanced lethal system in a single individual was performed in five runs of 10,000 replicates with identical default settings, across the 50,000 replicates we observed a total of 109 instances (e.g. Fig S7). of the balanced lethal system surviving until the end of simulation at generation 1,000.

**Table S7. Samples used in this study.** Species abbreviations: T. iva - *Triturus ivanbureschi*, T. mac - *Triturus macedonicus*, L. vul - *Lissotriton vulgaris*, L. mon - *Lissotriton montandoni*. ***** Two female *T. ivanbureschi* grandparents were captured at Zli Dol, Pčinja District, Serbia (42°25 N; 22°27 E). † Two male *T. macedonicus* grandparents were captured at Ceklin, Cetinje Municipality, Montenegro (42°21 N; 18°59 E). Ploidy analysis in this study uses raw sequences data from samples previously reported (*25*), all other samples are first reported in this study. *Triturus* chromosome 1 genotypes are reported as AB, AA and BB following the terminology of Macgregor & Horner (*4*).

| Sample | Taxon | Stage | Sex | Genotype | Purpose | Reference | SRA Run Accession |
|---|---|---|---|---|---|---|---|
| **BW_0012** | $F_1$ *T.iva* × *T.mac* | Adult | Male | AB | Trit map (father) | - | SRR31053713 |
| **BW_0013** | $F_1$ *T.iva* × *T.mac* | Adult | Female | AB | Trit map (mother) | - | SRR31053712 |
| **BW_0016*** | *T.iva* | Adult | Female | AB | Trit map (grandmother) | - | SRR31053575 |
| **BW_0017*** | *T.iva* | Adult | Female | AB | Trit map (grandmother) | - | SRR31053752 |
| **BW_0020†** | *T.mac* | Adult | Male | AB | Trit map (grandfather) | - | SRR31053619 |
| **BW_0022†** | *T.mac* | Adult | Male | AB | Trit map (grandfather) | - | SRR31053726 |
| **BW_0024** | $F_1$ *T.iva* × *T.mac* | Hatchling | unknown | AB | Ploidy | *(25)* | SRR29029578 |
| **BW_0025** | $F_1$ *T.iva* × *T.mac* | Hatchling | unknown | AB | Ploidy | *(25)* | SRR29029577 |
| **BW_0026** | $F_1$ *T.iva* × *T.mac* | Hatchling | unknown | AB | Ploidy | *(25)* | SRR29029566 |
| **BW_0027** | $F_1$ *T.iva* × *T.mac* | Hatchling | unknown | AB | Ploidy | *(25)* | SRR29029555 |
| **BW_0028** | $F_1$ *T.iva* × *T.mac* | Hatchling | unknown | AB | Ploidy | *(25)* | SRR29029554 |
| **BW_0029** | $F_1$ *T.iva* × *T.mac* | Hatchling | unknown | AB | Ploidy | *(25)* | SRR29029553 |
| **BW_0030** | $F_1$ *T.iva* × *T.mac* | Hatchling | unknown | AB | Ploidy | *(25)* | SRR29029552 |
| **BW_0031** | $F_1$ *T.iva* × *T.mac* | Hatchling | unknown | AB | Ploidy | *(25)* | SRR29029551 |
| **BW_0032** | $F_1$ *T.iva* × *T.mac* | Embryo | unknown | BB | Ploidy | *(25)* | SRR29029550 |
| **BW_0033** | $F_1$ *T.iva* × *T.mac* | Embryo | unknown | BB | Ploidy | *(25)* | SRR29029549 |
| **BW_0034** | $F_1$ *T.iva* × *T.mac* | Embryo | unknown | BB | Ploidy | *(25)* | SRR29029576 |
| **BW_0035** | $F_1$ *T.iva* × *T.mac* | Embryo | unknown | BB | Ploidy | *(25)* | SRR29029575 |
| **BW_0036** | $F_1$ *T.iva* × *T.mac* | Embryo | unknown | BB | Ploidy | *(25)* | SRR29029574 |
| **BW_0037** | $F_1$ *T.iva* × *T.mac* | Embryo | unknown | BB | Ploidy | *(25)* | SRR29029573 |
| **BW_0038** | $F_1$ *T.iva* × *T.mac* | Embryo | unknown | BB | Ploidy | *(25)* | SRR29029572 |
| **BW_0039** | $F_1$ *T.iva* × *T.mac* | Embryo | unknown | BB | Ploidy | *(25)* | SRR29029571 |
| **BW_0040** | $F_1$ *T.iva* × *T.mac* | Embryo | unknown | AA | Ploidy | *(25)* | SRR29029570 |
| **BW_0041** | $F_1$ *T.iva* × *T.mac* | Embryo | unknown | AA | Ploidy | *(25)* | SRR29029569 |
| **BW_0042** | $F_1$ *T.iva* × *T.mac* | Embryo | unknown | AA | Ploidy | *(25)* | SRR29029568 |
| **BW_0043** | $F_1$ *T.iva* × *T.mac* | Embryo | unknown | AA | Ploidy | *(25)* | SRR29029567 |
| **BW_0044** | $F_1$ *T.iva* × *T.mac* | Embryo | unknown | AA | Ploidy | *(25)* | SRR29029565 |
| **BW_0045** | $F_1$ *T.iva* × *T.mac* | Embryo | unknown | AA | Ploidy | *(25)* | SRR29029564 |
| **BW_0046** | $F_1$ *T.iva* × *T.mac* | Embryo | unknown | AA | Ploidy | *(25)* | SRR29029563 |
| **BW_0047** | $F_1$ *T.iva* × *T.mac* | Embryo | unknown | AA | Ploidy | *(25)* | SRR29029562 |
| **BW_0051** | $F_1$ *T.iva* × *T.mac* | Hatchling | unknown | AB | Ploidy | *(25)* | SRR29029561 |
| **BW_0052** | $F_1$ *T.iva* × *T.mac* | Hatchling | unknown | AB | Ploidy | *(25)* | SRR29029560 |
| **BW_0056** | $F_1$ *T.iva* × *T.mac* | Embryo | unknown | BB | Ploidy | *(25)* | SRR29029559 |
| **BW_0057** | $F_1$ *T.iva* × *T.mac* | Embryo | unknown | BB | Ploidy | *(25)* | SRR29029558 |
| **BW_0064** | $F_1$ *T.iva* × *T.mac* | Embryo | unknown | AA | Ploidy | *(25)* | SRR29029557 |
| **BW_0065** | $F_1$ *T.iva* × *T.mac* | Embryo | unknown | AA | Ploidy | *(25)* | SRR29029556 |
| **BW_0096** | $F_1$ *L.vul* × *L.mon* | Adult | Female | - | Liss map (father) | - | SRR31053888 |
| **BW_0097** | $F_1$ *L.vul* × *L.mon* | Adult | Male | - | Liss map (mother) | - | SRR31053887 |
| **BW_0098** | $F_2$ *T.iva* × *T.mac* | Hatchling | unknown | AB | Trit map (offspring) | - | SRR31053714 |
| **BW_0099** | $F_2$ *T.iva* × *T.mac* | Hatchling | unknown | AB | Trit map (offspring) | - | SRR31053505 |
| **BW_0100** | $F_2$ *T.iva* × *T.mac* | Hatchling | unknown | AB | Trit map (offspring) | - | SRR31053715 |
| **BW_0101** | $F_2$ *T.iva* × *T.mac* | Hatchling | unknown | AB | Trit map (offspring) | - | SRR31053811 |
| **BW_0102** | $F_2$ *T.iva* × *T.mac* | Hatchling | unknown | AB | Trit map (offspring) | - | SRR31053711 |
| **BW_0103** | $F_2$ *T.iva* × *T.mac* | Hatchling | unknown | AB | Trit map (offspring) | - | SRR31053700 |

| | | | | | | | |
|---|---|---|---|---|---|---|---|
| **BW_0104** | $F_2$ *T.iva* × *T.mac* | Hatchling | unknown | AB | Trit map (offspring) | - | SRR31053599 |
| **BW_0105** | $F_2$ *T.iva* × *T.mac* | Hatchling | unknown | AB | Trit map (offspring) | - | SRR31053592 |
| **BW_0106** | $F_2$ *T.iva* × *T.mac* | Hatchling | unknown | AB | Trit map (offspring) | - | SRR31053906 |
| **BW_0107** | $F_2$ *T.iva* × *T.mac* | Hatchling | unknown | AB | Trit map (offspring) | - | SRR31053894 |
| **BW_0108** | $F_2$ *T.iva* × *T.mac* | Hatchling | unknown | AB | Trit map (offspring) | - | SRR31053686 |
| **BW_0109** | $F_2$ *T.iva* × *T.mac* | Hatchling | unknown | AB | Trit map (offspring) | - | SRR31053676 |
| **BW_0110** | $F_2$ *T.iva* × *T.mac* | Hatchling | unknown | AB | Trit map (offspring) | - | SRR31053796 |
| **BW_0111** | $F_2$ *T.iva* × *T.mac* | Hatchling | unknown | AB | Trit map (offspring) | - | SRR31053784 |
| **BW_0112** | $F_2$ *T.iva* × *T.mac* | Hatchling | unknown | AB | Trit map (offspring) | - | SRR31053574 |
| **BW_0113** | $F_2$ *T.iva* × *T.mac* | Hatchling | unknown | AB | Trit map (offspring) | - | SRR31053563 |
| **BW_0114** | $F_2$ *T.iva* × *T.mac* | Hatchling | unknown | AB | Trit map (offspring) | - | SRR31053879 |
| **BW_0115** | $F_2$ *T.iva* × *T.mac* | Hatchling | unknown | AB | Trit map (offspring) | - | SRR31053884 |
| **BW_0116** | $F_2$ *T.iva* × *T.mac* | Hatchling | unknown | AB | Trit map (offspring) | - | SRR31053666 |
| **BW_0117** | $F_2$ *T.iva* × *T.mac* | Hatchling | unknown | AB | Trit map (offspring) | - | SRR31053652 |
| **BW_0122** | $F_2$ *T.iva* × *T.mac* | Hatchling | unknown | AB | Trit map (offspring) | - | SRR31053773 |
| **BW_0123** | $F_2$ *T.iva* × *T.mac* | Hatchling | unknown | AB | Trit map (offspring) | - | SRR31053762 |
| **BW_0124** | $F_2$ *T.iva* × *T.mac* | Hatchling | unknown | AB | Trit map (offspring) | - | SRR31053640 |
| **BW_0125** | $F_2$ *T.iva* × *T.mac* | Hatchling | unknown | AB | Trit map (offspring) | - | SRR31053629 |
| **BW_0126** | $F_2$ *T.iva* × *T.mac* | Hatchling | unknown | AB | Trit map (offspring) | - | SRR31053751 |
| **BW_0127** | $F_2$ *T.iva* × *T.mac* | Hatchling | unknown | AB | Trit map (offspring) | - | SRR31053740 |
| **BW_0128** | $F_2$ *T.iva* × *T.mac* | Hatchling | unknown | AB | Trit map (offspring) | - | SRR31053552 |
| **BW_0129** | $F_2$ *T.iva* × *T.mac* | Hatchling | unknown | AB | Trit map (offspring) | - | SRR31053541 |
| **BW_0130** | $F_2$ *T.iva* × *T.mac* | Hatchling | unknown | AB | Trit map (offspring) | - | SRR31053857 |
| **BW_0131** | $F_2$ *T.iva* × *T.mac* | Hatchling | unknown | AB | Trit map (offspring) | - | SRR31053846 |
| **BW_0132** | $F_2$ *T.iva* × *T.mac* | Hatchling | unknown | AB | Trit map (offspring) | - | SRR31053529 |
| **BW_0133** | $F_2$ *T.iva* × *T.mac* | Hatchling | unknown | AB | Trit map (offspring) | - | SRR31053518 |
| **BW_0134** | $F_2$ *T.iva* × *T.mac* | Hatchling | unknown | AB | Trit map (offspring) | - | SRR31053835 |
| **BW_0135** | $F_2$ *T.iva* × *T.mac* | Hatchling | unknown | AB | Trit map (offspring) | - | SRR31053824 |
| **BW_0136** | $F_2$ *T.iva* × *T.mac* | Hatchling | unknown | AB | Trit map (offspring) | - | SRR31053618 |
| **BW_0137** | $F_2$ *T.iva* × *T.mac* | Hatchling | unknown | AB | Trit map (offspring) | - | SRR31053607 |
| **BW_0138** | $F_2$ *T.iva* × *T.mac* | Hatchling | unknown | AB | Trit map (offspring) | - | SRR31053734 |
| **BW_0139** | $F_2$ *T.iva* × *T.mac* | Hatchling | unknown | AB | Trit map (offspring) | - | SRR31053733 |
| **BW_0140** | $F_2$ *T.iva* × *T.mac* | Hatchling | unknown | AB | Trit map (offspring) | - | SRR31053732 |
| **BW_0141** | $F_2$ *T.iva* × *T.mac* | Hatchling | unknown | AB | Trit map (offspring) | - | SRR31053731 |
| **BW_0142** | $F_2$ *T.iva* × *T.mac* | Hatchling | unknown | AB | Trit map (offspring) | - | SRR31053730 |
| **BW_0143** | $F_2$ *T.iva* × *T.mac* | Hatchling | unknown | AB | Trit map (offspring) | - | SRR31053729 |
| **BW_0145** | $F_2$ *T.iva* × *T.mac* | Hatchling | unknown | AB | Trit map (offspring) | - | SRR31053728 |
| **BW_0146** | $F_2$ *T.iva* × *T.mac* | Embryo | unknown | AA | Trit map (offspring) | - | SRR31053727 |
| **BW_0147** | $F_2$ *T.iva* × *T.mac* | Embryo | unknown | BB | Trit map (offspring) | - | SRR31053725 |
| **BW_0148** | $F_2$ *T.iva* × *T.mac* | Embryo | unknown | AB | Trit map (offspring) | - | SRR31053724 |
| **BW_0150** | $F_2$ *T.iva* × *T.mac* | Embryo | unknown | AA | Trit map (offspring) | - | SRR31053723 |
| **BW_0151** | $F_2$ *T.iva* × *T.mac* | Embryo | unknown | AA | Trit map (offspring) | - | SRR31053722 |
| **BW_0152** | $F_2$ *T.iva* × *T.mac* | Embryo | unknown | AA | Trit map (offspring) | - | SRR31053721 |
| **BW_0153** | $F_2$ *T.iva* × *T.mac* | Embryo | unknown | AA | Trit map (offspring) | - | SRR31053720 |
| **BW_0154** | $F_2$ *T.iva* × *T.mac* | Embryo | unknown | AA | Trit map (offspring) | - | SRR31053719 |
| **BW_0155** | $F_2$ *T.iva* × *T.mac* | Embryo | unknown | AA | Trit map (offspring) | - | SRR31053718 |
| **BW_0156** | $F_2$ *T.iva* × *T.mac* | Embryo | unknown | AA | Trit map (offspring) | - | SRR31053717 |
| **BW_0157** | $F_2$ *T.iva* × *T.mac* | Embryo | unknown | BB | Trit map (offspring) | - | SRR31053716 |
| **BW_0158** | $F_2$ *T.iva* × *T.mac* | Embryo | unknown | AA | Trit map (offspring) | - | SRR31053515 |
| **BW_0159** | $F_2$ *T.iva* × *T.mac* | Embryo | unknown | AB | Trit map (offspring) | - | SRR31053514 |
| **BW_0160** | $F_2$ *T.iva* × *T.mac* | Embryo | unknown | BB | Trit map (offspring) | - | SRR31053513 |
| **BW_0161** | $F_2$ *T.iva* × *T.mac* | Embryo | unknown | AB | Trit map (offspring) | - | SRR31053512 |
| **BW_0162** | $F_2$ *T.iva* × *T.mac* | Embryo | unknown | AA | Trit map (offspring) | - | SRR31053511 |
| **BW_0163** | $F_2$ *T.iva* × *T.mac* | Embryo | unknown | AB | Trit map (offspring) | - | SRR31053510 |
| **BW_0164** | $F_2$ *T.iva* × *T.mac* | Embryo | unknown | AA | Trit map (offspring) | - | SRR31053509 |
| **BW_0165** | $F_2$ *T.iva* × *T.mac* | Embryo | unknown | AA | Trit map (offspring) | - | SRR31053508 |

| | | | | | | | |
|---|---|---|---|---|---|---|---|
| **BW_0166** | $F_2$ *T.iva* × *T.mac* | Embryo | unknown | BB | Trit map (offspring) | - | SRR31053507 |
| **BW_0167** | $F_2$ *T.iva* × *T.mac* | Embryo | unknown | AA | Trit map (offspring) | - | SRR31053506 |
| **BW_0168** | $F_2$ *T.iva* × *T.mac* | Embryo | unknown | BB | Trit map (offspring) | - | SRR31053504 |
| **BW_0169** | $F_2$ *T.iva* × *T.mac* | Embryo | unknown | AA | Trit map (offspring) | - | SRR31053503 |
| **BW_0170** | $F_2$ *T.iva* × *T.mac* | Embryo | unknown | BB | Trit map (offspring) | - | SRR31053502 |
| **BW_0171** | $F_2$ *T.iva* × *T.mac* | Embryo | unknown | AB | Trit map (offspring) | - | SRR31053501 |
| **BW_0172** | $F_2$ *T.iva* × *T.mac* | Embryo | unknown | BB | Trit map (offspring) | - | SRR31053500 |
| **BW_0173** | $F_2$ *T.iva* × *T.mac* | Embryo | unknown | AA | Trit map (offspring) | - | SRR31053499 |
| **BW_0174** | $F_2$ *T.iva* × *T.mac* | Embryo | unknown | AB | Trit map (offspring) | - | SRR31053498 |
| **BW_0175** | $F_2$ *T.iva* × *T.mac* | Embryo | unknown | BB | Trit map (offspring) | - | SRR31053497 |
| **BW_0176** | $F_2$ *T.iva* × *T.mac* | Embryo | unknown | BB | Trit map (offspring) | - | SRR31053496 |
| **BW_0177** | $F_2$ *T.iva* × *T.mac* | Embryo | unknown | AA | Trit map (offspring) | - | SRR31053495 |
| **BW_0178** | $F_2$ *T.iva* × *T.mac* | Embryo | unknown | BB | Trit map (offspring) | - | SRR31053909 |
| **BW_0179** | $F_2$ *T.iva* × *T.mac* | Embryo | unknown | BB | Trit map (offspring) | - | SRR31053820 |
| **BW_0180** | $F_2$ *T.iva* × *T.mac* | Embryo | unknown | AB | Trit map (offspring) | - | SRR31053819 |
| **BW_0181** | $F_2$ *T.iva* × *T.mac* | Embryo | unknown | AB | Trit map (offspring) | - | SRR31053818 |
| **BW_0182** | $F_2$ *T.iva* × *T.mac* | Embryo | unknown | AA | Trit map (offspring) | - | SRR31053817 |
| **BW_0183** | $F_2$ *T.iva* × *T.mac* | Embryo | unknown | AB | Trit map (offspring) | - | SRR31053816 |
| **BW_0184** | $F_2$ *T.iva* × *T.mac* | Embryo | unknown | BB | Trit map (offspring) | - | SRR31053815 |
| **BW_0185** | $F_2$ *T.iva* × *T.mac* | Embryo | unknown | BB | Trit map (offspring) | - | SRR31053814 |
| **BW_0186** | $F_2$ *T.iva* × *T.mac* | Embryo | unknown | AA | Trit map (offspring) | - | SRR31053813 |
| **BW_0187** | $F_2$ *T.iva* × *T.mac* | Embryo | unknown | AA | Trit map (offspring) | - | SRR31053812 |
| **BW_0188** | $F_2$ *T.iva* × *T.mac* | Embryo | unknown | AA | Trit map (offspring) | - | SRR31053810 |
| **BW_0189** | $F_2$ *T.iva* × *T.mac* | Embryo | unknown | AA | Trit map (offspring) | - | SRR31053910 |
| **BW_0190** | $F_2$ *T.iva* × *T.mac* | Embryo | unknown | AA | Trit map (offspring) | - | SRR31053809 |
| **BW_0191** | $F_2$ *T.iva* × *T.mac* | Embryo | unknown | BB | Trit map (offspring) | - | SRR31053808 |
| **BW_0192** | $F_2$ *T.iva* × *T.mac* | Embryo | unknown | AA | Trit map (offspring) | - | SRR31053807 |
| **BW_0193** | $F_2$ *T.iva* × *T.mac* | Embryo | unknown | AA | Trit map (offspring) | - | SRR31053806 |
| **BW_0194** | $F_2$ *T.iva* × *T.mac* | Hatchling | unknown | AB | Trit map (offspring) | - | SRR31053805 |
| **BW_0195** | $F_2$ *T.iva* × *T.mac* | Hatchling | unknown | AB | Trit map (offspring) | - | SRR31053804 |
| **BW_0196** | $F_2$ *T.iva* × *T.mac* | Hatchling | unknown | AB | Trit map (offspring) | - | SRR31053803 |
| **BW_0197** | $F_2$ *T.iva* × *T.mac* | Hatchling | unknown | AB | Trit map (offspring) | - | SRR31053801 |
| **BW_0198** | $F_2$ *T.iva* × *T.mac* | Hatchling | unknown | AB | Trit map (offspring) | - | SRR31053710 |
| **BW_0199** | $F_2$ *T.iva* × *T.mac* | Hatchling | unknown | AB | Trit map (offspring) | - | SRR31053709 |
| **BW_0200** | $F_2$ *T.iva* × *T.mac* | Hatchling | unknown | AB | Trit map (offspring) | - | SRR31053708 |
| **BW_0201** | $F_2$ *T.iva* × *T.mac* | Hatchling | unknown | AB | Trit map (offspring) | - | SRR31053707 |
| **BW_0202** | $F_2$ *T.iva* × *T.mac* | Hatchling | unknown | AB | Trit map (offspring) | - | SRR31053706 |
| **BW_0203** | $F_2$ *T.iva* × *T.mac* | Hatchling | unknown | AB | Trit map (offspring) | - | SRR31053705 |
| **BW_0204** | $F_2$ *T.iva* × *T.mac* | Hatchling | unknown | AB | Trit map (offspring) | - | SRR31053704 |
| **BW_0205** | $F_2$ *T.iva* × *T.mac* | Hatchling | unknown | AB | Trit map (offspring) | - | SRR31053703 |
| **BW_0206** | $F_2$ *T.iva* × *T.mac* | Hatchling | unknown | AB | Trit map (offspring) | - | SRR31053702 |
| **BW_0207** | $F_2$ *T.iva* × *T.mac* | Hatchling | unknown | AB | Trit map (offspring) | - | SRR31053701 |
| **BW_0208** | $F_2$ *T.iva* × *T.mac* | Hatchling | unknown | AB | Trit map (offspring) | - | SRR31053699 |
| **BW_0209** | $F_2$ *T.iva* × *T.mac* | Hatchling | unknown | AB | Trit map (offspring) | - | SRR31053698 |
| **BW_0210** | $F_2$ *T.iva* × *T.mac* | Hatchling | unknown | AB | Trit map (offspring) | - | SRR31053696 |
| **BW_0211** | $F_2$ *T.iva* × *T.mac* | Hatchling | unknown | AB | Trit map (offspring) | - | SRR31053695 |
| **BW_0212** | $F_2$ *T.iva* × *T.mac* | Hatchling | unknown | AB | Trit map (offspring) | - | SRR31053694 |
| **BW_0213** | $F_2$ *T.iva* × *T.mac* | Hatchling | unknown | AB | Trit map (offspring) | - | SRR31053697 |
| **BW_0214** | $F_2$ *T.iva* × *T.mac* | Hatchling | unknown | AB | Trit map (offspring) | - | SRR31053693 |
| **BW_0215** | $F_2$ *T.iva* × *T.mac* | Hatchling | unknown | AB | Trit map (offspring) | - | SRR31053692 |
| **BW_0216** | $F_2$ *T.iva* × *T.mac* | Hatchling | unknown | AB | Trit map (offspring) | - | SRR31053605 |
| **BW_0217** | $F_2$ *T.iva* × *T.mac* | Hatchling | unknown | AB | Trit map (offspring) | - | SRR31053601 |
| **BW_0218** | $F_2$ *T.iva* × *T.mac* | Hatchling | unknown | AB | Trit map (offspring) | - | SRR31053598 |
| **BW_0219** | $F_2$ *T.iva* × *T.mac* | Hatchling | unknown | AB | Trit map (offspring) | - | SRR31053597 |
| **BW_0220** | $F_2$ *T.iva* × *T.mac* | Hatchling | unknown | AB | Trit map (offspring) | - | SRR31053600 |
| **BW_0221** | $F_2$ *T.iva* × *T.mac* | Hatchling | unknown | AB | Trit map (offspring) | - | SRR31053596 |

| | | | | | | | |
|---|---|---|---|---|---|---|---|
| **BW_0222** | $F_2$ *T.iva* × *T.mac* | Hatchling | unknown | AB | Trit map (offspring) | - | SRR31053595 |
| **BW_0223** | $F_2$ *T.iva* × *T.mac* | Hatchling | unknown | AB | Trit map (offspring) | - | SRR31053594 |
| **BW_0224** | $F_2$ *T.iva* × *T.mac* | Embryo | unknown | AA | Trit map (offspring) | - | SRR31053593 |
| **BW_0225** | $F_2$ *T.iva* × *T.mac* | Embryo | unknown | AA | Trit map (offspring) | - | SRR31053591 |
| **BW_0226** | $F_2$ *T.iva* × *T.mac* | Embryo | unknown | AB | Trit map (offspring) | - | SRR31053590 |
| **BW_0227** | $F_2$ *T.iva* × *T.mac* | Embryo | unknown | AA | Trit map (offspring) | - | SRR31053589 |
| **BW_0228** | $F_2$ *T.iva* × *T.mac* | Embryo | unknown | AA | Trit map (offspring) | - | SRR31053588 |
| **BW_0229** | $F_2$ *T.iva* × *T.mac* | Embryo | unknown | AA | Trit map (offspring) | - | SRR31053587 |
| **BW_0230** | $F_2$ *T.iva* × *T.mac* | Embryo | unknown | BB | Trit map (offspring) | - | SRR31053586 |
| **BW_0231** | $F_2$ *T.iva* × *T.mac* | Embryo | unknown | AA | Trit map (offspring) | - | SRR31053585 |
| **BW_0232** | $F_2$ *T.iva* × *T.mac* | Embryo | unknown | AA | Trit map (offspring) | - | SRR31053584 |
| **BW_0233** | $F_2$ *T.iva* × *T.mac* | Embryo | unknown | AB | Trit map (offspring) | - | SRR31053583 |
| **BW_0235** | $F_2$ *T.iva* × *T.mac* | Embryo | unknown | AA | Trit map (offspring) | - | SRR31053582 |
| **BW_0236** | $F_2$ *T.iva* × *T.mac* | Embryo | unknown | AA | Trit map (offspring) | - | SRR31053911 |
| **BW_0237** | $F_2$ *T.iva* × *T.mac* | Embryo | unknown | BB | Trit map (offspring) | - | SRR31053908 |
| **BW_0238** | $F_2$ *T.iva* × *T.mac* | Embryo | unknown | BB | Trit map (offspring) | - | SRR31053907 |
| **BW_0239** | $F_2$ *T.iva* × *T.mac* | Embryo | unknown | BB | Trit map (offspring) | - | SRR31053905 |
| **BW_0240** | $F_2$ *T.iva* × *T.mac* | Embryo | unknown | AA | Trit map (offspring) | - | SRR31053904 |
| **BW_0241** | $F_2$ *T.iva* × *T.mac* | Embryo | unknown | AA | Trit map (offspring) | - | SRR31053903 |
| **BW_0242** | $F_2$ *T.iva* × *T.mac* | Embryo | unknown | AA | Trit map (offspring) | - | SRR31053902 |
| **BW_0243** | $F_2$ *T.iva* × *T.mac* | Embryo | unknown | AA | Trit map (offspring) | - | SRR31053901 |
| **BW_0244** | $F_2$ *T.iva* × *T.mac* | Embryo | unknown | AA | Trit map (offspring) | - | SRR31053900 |
| **BW_0245** | $F_2$ *T.iva* × *T.mac* | Embryo | unknown | AA | Trit map (offspring) | - | SRR31053899 |
| **BW_0246** | $F_2$ *T.iva* × *T.mac* | Embryo | unknown | AB | Trit map (offspring) | - | SRR31053898 |
| **BW_0247** | $F_2$ *T.iva* × *T.mac* | Embryo | unknown | BB | Trit map (offspring) | - | SRR31053896 |
| **BW_0248** | $F_2$ *T.iva* × *T.mac* | Embryo | unknown | BB | Trit map (offspring) | - | SRR31053895 |
| **BW_0249** | $F_2$ *T.iva* × *T.mac* | Embryo | unknown | AA | Trit map (offspring) | - | SRR31053897 |
| **BW_0250** | $F_2$ *T.iva* × *T.mac* | Embryo | unknown | BB | Trit map (offspring) | - | SRR31053893 |
| **BW_0251** | $F_2$ *T.iva* × *T.mac* | Embryo | unknown | BB | Trit map (offspring) | - | SRR31053892 |
| **BW_0252** | $F_2$ *T.iva* × *T.mac* | Embryo | unknown | BB | Trit map (offspring) | - | SRR31053891 |
| **BW_0253** | $F_2$ *T.iva* × *T.mac* | Embryo | unknown | BB | Trit map (offspring) | - | SRR31053890 |
| **BW_0254** | $F_2$ *T.iva* × *T.mac* | Embryo | unknown | BB | Trit map (offspring) | - | SRR31053889 |
| **BW_0255** | $F_2$ *T.iva* × *T.mac* | Embryo | unknown | AA | Trit map (offspring) | - | SRR31053691 |
| **BW_0256** | $F_2$ *T.iva* × *T.mac* | Embryo | unknown | BB | Trit map (offspring) | - | SRR31053690 |
| **BW_0257** | $F_2$ *T.iva* × *T.mac* | Embryo | unknown | AA | Trit map (offspring) | - | SRR31053688 |
| **BW_0258** | $F_2$ *T.iva* × *T.mac* | Embryo | unknown | BB | Trit map (offspring) | - | SRR31053687 |
| **BW_0259** | $F_2$ *T.iva* × *T.mac* | Embryo | unknown | AA | Trit map (offspring) | - | SRR31053689 |
| **BW_0262** | $F_2$ *T.iva* × *T.mac* | Embryo | unknown | BB | Trit map (offspring) | - | SRR31053685 |
| **BW_0263** | $F_2$ *T.iva* × *T.mac* | Embryo | unknown | AA | Trit map (offspring) | - | SRR31053684 |
| **BW_0264** | $F_2$ *T.iva* × *T.mac* | Embryo | unknown | BB | Trit map (offspring) | - | SRR31053683 |
| **BW_0266** | $F_2$ *T.iva* × *T.mac* | Embryo | unknown | AB | Trit map (offspring) | - | SRR31053682 |
| **BW_0267** | $F_2$ *T.iva* × *T.mac* | Embryo | unknown | AA | Trit map (offspring) | - | SRR31053680 |
| **BW_0270** | $F_2$ *T.iva* × *T.mac* | Embryo | unknown | AB | Trit map (offspring) | - | SRR31053679 |
| **BW_0271** | $F_2$ *T.iva* × *T.mac* | Embryo | unknown | AA | Trit map (offspring) | - | SRR31053678 |
| **BW_0272** | $F_2$ *T.iva* × *T.mac* | Embryo | unknown | AB | Trit map (offspring) | - | SRR31053681 |
| **BW_0273** | $F_2$ *T.iva* × *T.mac* | Embryo | unknown | AA | Trit map (offspring) | - | SRR31053677 |
| **BW_0274** | $F_2$ *T.iva* × *T.mac* | Embryo | unknown | AA | Trit map (offspring) | - | SRR31053675 |
| **BW_0275** | $F_2$ *T.iva* × *T.mac* | Embryo | unknown | BB | Trit map (offspring) | - | SRR31053674 |
| **BW_0278** | $F_2$ *T.iva* × *T.mac* | Embryo | unknown | AA | Trit map (offspring) | - | SRR31053673 |
| **BW_0279** | $F_2$ *T.iva* × *T.mac* | Embryo | unknown | AA | Trit map (offspring) | - | SRR31053672 |
| **BW_0280** | $F_2$ *T.iva* × *T.mac* | Embryo | unknown | AA | Trit map (offspring) | - | SRR31053671 |
| **BW_0281** | $F_2$ *T.iva* × *T.mac* | Embryo | unknown | BB | Trit map (offspring) | - | SRR31053802 |
| **BW_0282** | $F_2$ *T.iva* × *T.mac* | Embryo | unknown | AA | Trit map (offspring) | - | SRR31053800 |
| **BW_0283** | $F_2$ *T.iva* × *T.mac* | Embryo | unknown | BB | Trit map (offspring) | - | SRR31053799 |
| **BW_0284** | $F_2$ *T.iva* × *T.mac* | Embryo | unknown | BB | Trit map (offspring) | - | SRR31053798 |
| **BW_0285** | $F_2$ *T.iva* × *T.mac* | Embryo | unknown | BB | Trit map (offspring) | - | SRR31053797 |

| | | | | | | | |
|---|---|---|---|---|---|---|---|
| **BW_0286** | $F_2$ *T.iva* × *T.mac* | Embryo | unknown | BB | Trit map (offspring) | - | SRR31053795 |
| **BW_0287** | $F_2$ *T.iva* × *T.mac* | Embryo | unknown | BB | Trit map (offspring) | - | SRR31053794 |
| **BW_0288** | $F_2$ *T.iva* × *T.mac* | Embryo | unknown | AA | Trit map (offspring) | - | SRR31053793 |
| **BW_0289** | $F_2$ *T.iva* × *T.mac* | Embryo | unknown | AA | Trit map (offspring) | - | SRR31053792 |
| **BW_0290** | $F_2$ *T.iva* × *T.mac* | Embryo | unknown | BB | Trit map (offspring) | - | SRR31053791 |
| **BW_0291** | $F_2$ *T.iva* × *T.mac* | Embryo | unknown | BB | Trit map (offspring) | - | SRR31053790 |
| **BW_0292** | $F_2$ *T.iva* × *T.mac* | Embryo | unknown | AA | Trit map (offspring) | - | SRR31053789 |
| **BW_0293** | $F_2$ *T.iva* × *T.mac* | Embryo | unknown | AA | Trit map (offspring) | - | SRR31053788 |
| **BW_0294** | $F_2$ *T.iva* × *T.mac* | Embryo | unknown | AA | Trit map (offspring) | - | SRR31053787 |
| **BW_0295** | $F_2$ *T.iva* × *T.mac* | Embryo | unknown | AA | Trit map (offspring) | - | SRR31053786 |
| **BW_0296** | $F_2$ *T.iva* × *T.mac* | Embryo | unknown | BB | Trit map (offspring) | - | SRR31053783 |
| **BW_0297** | $F_2$ *T.iva* × *T.mac* | Embryo | unknown | AA | Trit map (offspring) | - | SRR31053782 |
| **BW_0298** | $F_2$ *T.iva* × *T.mac* | Embryo | unknown | BB | Trit map (offspring) | - | SRR31053781 |
| **BW_0299** | $F_2$ *T.iva* × *T.mac* | Embryo | unknown | BB | Trit map (offspring) | - | SRR31053581 |
| **BW_0300** | $F_2$ *T.iva* × *T.mac* | Embryo | unknown | AA | Trit map (offspring) | - | SRR31053580 |
| **BW_0301** | $F_2$ *T.iva* × *T.mac* | Embryo | unknown | BB | Trit map (offspring) | - | SRR31053579 |
| **BW_0302** | $F_2$ *T.iva* × *T.mac* | Embryo | unknown | BB | Trit map (offspring) | - | SRR31053578 |
| **BW_0303** | $F_2$ *T.iva* × *T.mac* | Embryo | unknown | BB | Trit map (offspring) | - | SRR31053577 |
| **BW_0304** | $F_2$ *T.iva* × *T.mac* | Embryo | unknown | BB | Trit map (offspring) | - | SRR31053785 |
| **BW_0305** | $F_2$ *T.iva* × *T.mac* | Embryo | unknown | BB | Trit map (offspring) | - | SRR31053576 |
| **BW_0306** | $F_2$ *T.iva* × *T.mac* | Embryo | unknown | AA | Trit map (offspring) | - | SRR31053573 |
| **BW_0307** | $F_2$ *T.iva* × *T.mac* | Embryo | unknown | BB | Trit map (offspring) | - | SRR31053572 |
| **BW_0308** | $F_2$ *T.iva* × *T.mac* | Embryo | unknown | BB | Trit map (offspring) | - | SRR31053571 |
| **BW_0309** | $F_2$ *T.iva* × *T.mac* | Embryo | unknown | BB | Trit map (offspring) | - | SRR31053570 |
| **BW_0310** | $F_2$ *T.iva* × *T.mac* | Embryo | unknown | BB | Trit map (offspring) | - | SRR31053568 |
| **BW_0311** | $F_2$ *T.iva* × *T.mac* | Embryo | unknown | AA | Trit map (offspring) | - | SRR31053567 |
| **BW_0313** | $F_2$ *T.iva* × *T.mac* | Embryo | unknown | BB | Trit map (offspring) | - | SRR31053566 |
| **BW_0314** | $F_2$ *T.iva* × *T.mac* | Embryo | unknown | BB | Trit map (offspring) | - | SRR31053569 |
| **BW_0315** | $F_2$ *T.iva* × *T.mac* | Embryo | unknown | BB | Trit map (offspring) | - | SRR31053565 |
| **BW_0316** | $F_2$ *T.iva* × *T.mac* | Embryo | unknown | BB | Trit map (offspring) | - | SRR31053564 |
| **BW_0318** | $F_2$ *T.iva* × *T.mac* | Embryo | unknown | AA | Trit map (offspring) | - | SRR31053562 |
| **BW_0319** | $F_2$ *T.iva* × *T.mac* | Embryo | unknown | BB | Trit map (offspring) | - | SRR31053560 |
| **BW_0321** | $F_2$ *L.vul* × *L.mon* | Hatchling | unknown | - | Liss map (offspring) | - | SRR31053886 |
| **BW_0322** | $F_2$ *L.vul* × *L.mon* | Hatchling | unknown | - | Liss map (offspring) | - | SRR31053885 |
| **BW_0323** | $F_2$ *L.vul* × *L.mon* | Hatchling | unknown | - | Liss map (offspring) | - | SRR31053883 |
| **BW_0324** | $F_2$ *L.vul* × *L.mon* | Hatchling | unknown | - | Liss map (offspring) | - | SRR31053882 |
| **BW_0325** | $F_2$ *L.vul* × *L.mon* | Hatchling | unknown | - | Liss map (offspring) | - | SRR31053881 |
| **BW_0326** | $F_2$ *L.vul* × *L.mon* | Hatchling | unknown | - | Liss map (offspring) | - | SRR31053880 |
| **BW_0327** | $F_2$ *L.vul* × *L.mon* | Hatchling | unknown | - | Liss map (offspring) | - | SRR31053878 |
| **BW_0329** | $F_2$ *L.vul* × *L.mon* | Hatchling | unknown | - | Liss map (offspring) | - | SRR31053877 |
| **BW_0330** | $F_2$ *L.vul* × *L.mon* | Hatchling | unknown | - | Liss map (offspring) | - | SRR31053876 |
| **BW_0331** | $F_2$ *L.vul* × *L.mon* | Hatchling | unknown | - | Liss map (offspring) | - | SRR31053875 |
| **BW_0332** | $F_2$ *L.vul* × *L.mon* | Hatchling | unknown | - | Liss map (offspring) | - | SRR31053874 |
| **BW_0333** | $F_2$ *L.vul* × *L.mon* | Hatchling | unknown | - | Liss map (offspring) | - | SRR31053873 |
| **BW_0334** | $F_2$ *L.vul* × *L.mon* | Hatchling | unknown | - | Liss map (offspring) | - | SRR31053872 |
| **BW_0337** | $F_2$ *L.vul* × *L.mon* | Hatchling | unknown | - | Liss map (offspring) | - | SRR31053871 |
| **BW_0338** | $F_2$ *L.vul* × *L.mon* | Hatchling | unknown | - | Liss map (offspring) | - | SRR31053870 |
| **BW_0339** | $F_2$ *L.vul* × *L.mon* | Hatchling | unknown | - | Liss map (offspring) | - | SRR31053869 |
| **BW_0340** | $F_2$ *L.vul* × *L.mon* | Hatchling | unknown | - | Liss map (offspring) | - | SRR31053868 |
| **BW_0341** | $F_2$ *L.vul* × *L.mon* | Hatchling | unknown | - | Liss map (offspring) | - | SRR31053867 |
| **BW_0342** | $F_2$ *L.vul* × *L.mon* | Hatchling | unknown | - | Liss map (offspring) | - | SRR31053866 |
| **BW_0343** | $F_2$ *L.vul* × *L.mon* | Hatchling | unknown | - | Liss map (offspring) | - | SRR31053670 |
| **BW_0344** | $F_2$ *L.vul* × *L.mon* | Hatchling | unknown | - | Liss map (offspring) | - | SRR31053669 |
| **BW_0345** | $F_2$ *L.vul* × *L.mon* | Hatchling | unknown | - | Liss map (offspring) | - | SRR31053668 |
| **BW_0346** | $F_2$ *L.vul* × *L.mon* | Hatchling | unknown | - | Liss map (offspring) | - | SRR31053667 |
| **BW_0347** | $F_2$ *L.vul* × *L.mon* | Hatchling | unknown | - | Liss map (offspring) | - | SRR31053665 |

| | | | | | | | |
|---|---|---|---|---|---|---|---|
| **BW_0348** | $F_2$ *L.vul* × *L.mon* | Hatchling | unknown | - | Liss map (offspring) | - | SRR31053664 |
| **BW_0349** | $F_2$ *L.vul* × *L.mon* | Hatchling | unknown | - | Liss map (offspring) | - | SRR31053663 |
| **BW_0350** | $F_2$ *L.vul* × *L.mon* | Hatchling | unknown | - | Liss map (offspring) | - | SRR31053662 |
| **BW_0351** | $F_2$ *L.vul* × *L.mon* | Hatchling | unknown | - | Liss map (offspring) | - | SRR31053661 |
| **BW_0353** | $F_2$ *L.vul* × *L.mon* | Hatchling | unknown | - | Liss map (offspring) | - | SRR31053660 |
| **BW_0354** | $F_2$ *L.vul* × *L.mon* | Hatchling | unknown | - | Liss map (offspring) | - | SRR31053659 |
| **BW_0355** | $F_2$ *L.vul* × *L.mon* | Hatchling | unknown | - | Liss map (offspring) | - | SRR31053657 |
| **BW_0356** | $F_2$ *L.vul* × *L.mon* | Hatchling | unknown | - | Liss map (offspring) | - | SRR31053656 |
| **BW_0357** | $F_2$ *L.vul* × *L.mon* | Hatchling | unknown | - | Liss map (offspring) | - | SRR31053655 |
| **BW_0358** | $F_2$ *L.vul* × *L.mon* | Hatchling | unknown | - | Liss map (offspring) | - | SRR31053658 |
| **BW_0359** | $F_2$ *L.vul* × *L.mon* | Hatchling | unknown | - | Liss map (offspring) | - | SRR31053654 |
| **BW_0360** | $F_2$ *L.vul* × *L.mon* | Hatchling | unknown | - | Liss map (offspring) | - | SRR31053653 |
| **BW_0362** | $F_2$ *L.vul* × *L.mon* | Hatchling | unknown | - | Liss map (offspring) | - | SRR31053651 |
| **BW_0363** | $F_2$ *L.vul* × *L.mon* | Hatchling | unknown | - | Liss map (offspring) | - | SRR31053649 |
| **BW_0364** | $F_2$ *L.vul* × *L.mon* | Hatchling | unknown | - | Liss map (offspring) | - | SRR31053780 |
| **BW_0365** | $F_2$ *L.vul* × *L.mon* | Hatchling | unknown | - | Liss map (offspring) | - | SRR31053779 |
| **BW_0366** | $F_2$ *L.vul* × *L.mon* | Hatchling | unknown | - | Liss map (offspring) | - | SRR31053650 |
| **BW_0367** | $F_2$ *L.vul* × *L.mon* | Hatchling | unknown | - | Liss map (offspring) | - | SRR31053778 |
| **BW_0368** | $F_2$ *L.vul* × *L.mon* | Hatchling | unknown | - | Liss map (offspring) | - | SRR31053777 |
| **BW_0369** | $F_2$ *L.vul* × *L.mon* | Hatchling | unknown | - | Liss map (offspring) | - | SRR31053776 |
| **BW_0370** | $F_2$ *L.vul* × *L.mon* | Hatchling | unknown | - | Liss map (offspring) | - | SRR31053775 |
| **BW_0371** | $F_2$ *L.vul* × *L.mon* | Hatchling | unknown | - | Liss map (offspring) | - | SRR31053774 |
| **BW_0372** | $F_2$ *L.vul* × *L.mon* | Hatchling | unknown | - | Liss map (offspring) | - | SRR31053772 |
| **BW_0373** | $F_2$ *L.vul* × *L.mon* | Hatchling | unknown | - | Liss map (offspring) | - | SRR31053771 |
| **BW_0374** | $F_2$ *L.vul* × *L.mon* | Hatchling | unknown | - | Liss map (offspring) | - | SRR31053770 |
| **BW_0375** | $F_2$ *L.vul* × *L.mon* | Hatchling | unknown | - | Liss map (offspring) | - | SRR31053769 |
| **BW_0376** | $F_2$ *L.vul* × *L.mon* | Hatchling | unknown | - | Liss map (offspring) | - | SRR31053768 |
| **BW_0377** | $F_2$ *L.vul* × *L.mon* | Hatchling | unknown | - | Liss map (offspring) | - | SRR31053767 |
| **BW_0378** | $F_2$ *L.vul* × *L.mon* | Hatchling | unknown | - | Liss map (offspring) | - | SRR31053765 |
| **BW_0379** | $F_2$ *L.vul* × *L.mon* | Hatchling | unknown | - | Liss map (offspring) | - | SRR31053764 |
| **BW_0381** | $F_2$ *L.vul* × *L.mon* | Hatchling | unknown | - | Liss map (offspring) | - | SRR31053763 |
| **BW_0382** | $F_2$ *L.vul* × *L.mon* | Hatchling | unknown | - | Liss map (offspring) | - | SRR31053766 |
| **BW_0383** | $F_2$ *L.vul* × *L.mon* | Hatchling | unknown | - | Liss map (offspring) | - | SRR31053761 |
| **BW_0384** | $F_2$ *L.vul* × *L.mon* | Hatchling | unknown | - | Liss map (offspring) | - | SRR31053759 |
| **BW_0385** | $F_2$ *L.vul* × *L.mon* | Hatchling | unknown | - | Liss map (offspring) | - | SRR31053648 |
| **BW_0386** | $F_2$ *L.vul* × *L.mon* | Hatchling | unknown | - | Liss map (offspring) | - | SRR31053647 |
| **BW_0387** | $F_2$ *L.vul* × *L.mon* | Hatchling | unknown | - | Liss map (offspring) | - | SRR31053646 |
| **BW_0388** | $F_2$ *L.vul* × *L.mon* | Hatchling | unknown | - | Liss map (offspring) | - | SRR31053645 |
| **BW_0389** | $F_2$ *L.vul* × *L.mon* | Hatchling | unknown | - | Liss map (offspring) | - | SRR31053644 |
| **BW_0390** | $F_2$ *L.vul* × *L.mon* | Hatchling | unknown | - | Liss map (offspring) | - | SRR31053643 |
| **BW_0391** | $F_2$ *L.vul* × *L.mon* | Hatchling | unknown | - | Liss map (offspring) | - | SRR31053642 |
| **BW_0392** | $F_2$ *L.vul* × *L.mon* | Hatchling | unknown | - | Liss map (offspring) | - | SRR31053641 |
| **BW_0393** | $F_2$ *L.vul* × *L.mon* | Hatchling | unknown | - | Liss map (offspring) | - | SRR31053639 |
| **BW_0394** | $F_2$ *L.vul* × *L.mon* | Hatchling | unknown | - | Liss map (offspring) | - | SRR31053638 |
| **BW_0396** | $F_2$ *L.vul* × *L.mon* | Hatchling | unknown | - | Liss map (offspring) | - | SRR31053637 |
| **BW_0397** | $F_2$ *L.vul* × *L.mon* | Hatchling | unknown | - | Liss map (offspring) | - | SRR31053636 |
| **BW_0398** | $F_2$ *L.vul* × *L.mon* | Hatchling | unknown | - | Liss map (offspring) | - | SRR31053635 |
| **BW_0399** | $F_2$ *L.vul* × *L.mon* | Hatchling | unknown | - | Liss map (offspring) | - | SRR31053634 |
| **BW_0400** | $F_2$ *L.vul* × *L.mon* | Hatchling | unknown | - | Liss map (offspring) | - | SRR31053633 |
| **BW_0401** | $F_2$ *L.vul* × *L.mon* | Hatchling | unknown | - | Liss map (offspring) | - | SRR31053632 |
| **BW_0402** | $F_2$ *L.vul* × *L.mon* | Hatchling | unknown | - | Liss map (offspring) | - | SRR31053631 |
| **BW_0403** | $F_2$ *L.vul* × *L.mon* | Hatchling | unknown | - | Liss map (offspring) | - | SRR31053630 |
| **BW_0404** | $F_2$ *L.vul* × *L.mon* | Hatchling | unknown | - | Liss map (offspring) | - | SRR31053628 |
| **BW_0405** | $F_2$ *L.vul* × *L.mon* | Hatchling | unknown | - | Liss map (offspring) | - | SRR31053627 |
| **BW_0406** | $F_2$ *L.vul* × *L.mon* | Hatchling | unknown | - | Liss map (offspring) | - | SRR31053626 |
| **BW_0407** | $F_2$ *L.vul* × *L.mon* | Hatchling | unknown | - | Liss map (offspring) | - | SRR31053760 |

| | | | | | | | |
|---|---|---|---|---|---|---|---|
| **BW_0408** | $F_2$ *L.vul* × *L.mon* | Hatchling | unknown | - | Liss map (offspring) | - | SRR31053758 |
| **BW_0409** | $F_2$ *L.vul* × *L.mon* | Hatchling | unknown | - | Liss map (offspring) | - | SRR31053756 |
| **BW_0410** | $F_2$ *L.vul* × *L.mon* | Hatchling | unknown | - | Liss map (offspring) | - | SRR31053755 |
| **BW_0411** | $F_2$ *L.vul* × *L.mon* | Hatchling | unknown | - | Liss map (offspring) | - | SRR31053754 |
| **BW_0412** | $F_2$ *L.vul* × *L.mon* | Hatchling | unknown | - | Liss map (offspring) | - | SRR31053757 |
| **BW_0413** | $F_2$ *L.vul* × *L.mon* | Hatchling | unknown | - | Liss map (offspring) | - | SRR31053753 |
| **BW_0414** | $F_2$ *L.vul* × *L.mon* | Hatchling | unknown | - | Liss map (offspring) | - | SRR31053750 |
| **BW_0415** | $F_2$ *L.vul* × *L.mon* | Hatchling | unknown | - | Liss map (offspring) | - | SRR31053749 |
| **BW_0416** | $F_2$ *L.vul* × *L.mon* | Hatchling | unknown | - | Liss map (offspring) | - | SRR31053748 |
| **BW_0417** | $F_2$ *L.vul* × *L.mon* | Hatchling | unknown | - | Liss map (offspring) | - | SRR31053747 |
| **BW_0418** | $F_2$ *L.vul* × *L.mon* | Hatchling | unknown | - | Liss map (offspring) | - | SRR31053745 |
| **BW_0419** | $F_2$ *L.vul* × *L.mon* | Hatchling | unknown | - | Liss map (offspring) | - | SRR31053744 |
| **BW_0420** | $F_2$ *L.vul* × *L.mon* | Hatchling | unknown | - | Liss map (offspring) | - | SRR31053743 |
| **BW_0421** | $F_2$ *L.vul* × *L.mon* | Hatchling | unknown | - | Liss map (offspring) | - | SRR31053746 |
| **BW_0422** | $F_2$ *L.vul* × *L.mon* | Hatchling | unknown | - | Liss map (offspring) | - | SRR31053742 |
| **BW_0423** | $F_2$ *L.vul* × *L.mon* | Hatchling | unknown | - | Liss map (offspring) | - | SRR31053741 |
| **BW_0424** | $F_2$ *L.vul* × *L.mon* | Hatchling | unknown | - | Liss map (offspring) | - | SRR31053739 |
| **BW_0425** | $F_2$ *L.vul* × *L.mon* | Hatchling | unknown | - | Liss map (offspring) | - | SRR31053738 |
| **BW_0426** | $F_2$ *L.vul* × *L.mon* | Hatchling | unknown | - | Liss map (offspring) | - | SRR31053561 |
| **BW_0427** | $F_2$ *L.vul* × *L.mon* | Hatchling | unknown | - | Liss map (offspring) | - | SRR31053559 |
| **BW_0428** | $F_2$ *L.vul* × *L.mon* | Hatchling | unknown | - | Liss map (offspring) | - | SRR31053557 |
| **BW_0429** | $F_2$ *L.vul* × *L.mon* | Hatchling | unknown | - | Liss map (offspring) | - | SRR31053556 |
| **BW_0430** | $F_2$ *L.vul* × *L.mon* | Hatchling | unknown | - | Liss map (offspring) | - | SRR31053555 |
| **BW_0431** | $F_2$ *L.vul* × *L.mon* | Hatchling | unknown | - | Liss map (offspring) | - | SRR31053558 |
| **BW_0432** | $F_2$ *L.vul* × *L.mon* | Hatchling | unknown | - | Liss map (offspring) | - | SRR31053554 |
| **BW_0433** | $F_2$ *L.vul* × *L.mon* | Hatchling | unknown | - | Liss map (offspring) | - | SRR31053553 |
| **BW_0434** | $F_2$ *L.vul* × *L.mon* | Hatchling | unknown | - | Liss map (offspring) | - | SRR31053551 |
| **BW_0435** | $F_2$ *L.vul* × *L.mon* | Hatchling | unknown | - | Liss map (offspring) | - | SRR31053550 |
| **BW_0436** | $F_2$ *L.vul* × *L.mon* | Hatchling | unknown | - | Liss map (offspring) | - | SRR31053549 |
| **BW_0438** | $F_2$ *L.vul* × *L.mon* | Hatchling | unknown | - | Liss map (offspring) | - | SRR31053548 |
| **BW_0439** | $F_2$ *L.vul* × *L.mon* | Hatchling | unknown | - | Liss map (offspring) | - | SRR31053546 |
| **BW_0440** | $F_2$ *L.vul* × *L.mon* | Hatchling | unknown | - | Liss map (offspring) | - | SRR31053545 |
| **BW_0441** | $F_2$ *L.vul* × *L.mon* | Hatchling | unknown | - | Liss map (offspring) | - | SRR31053544 |
| **BW_0442** | $F_2$ *L.vul* × *L.mon* | Hatchling | unknown | - | Liss map (offspring) | - | SRR31053547 |
| **BW_0443** | $F_2$ *L.vul* × *L.mon* | Hatchling | unknown | - | Liss map (offspring) | - | SRR31053543 |
| **BW_0444** | $F_2$ *L.vul* × *L.mon* | Hatchling | unknown | - | Liss map (offspring) | - | SRR31053542 |
| **BW_0446** | $F_2$ *L.vul* × *L.mon* | Hatchling | unknown | - | Liss map (offspring) | - | SRR31053540 |
| **BW_0447** | $F_2$ *L.vul* × *L.mon* | Hatchling | unknown | - | Liss map (offspring) | - | SRR31053538 |
| **BW_0448** | $F_2$ *L.vul* × *L.mon* | Hatchling | unknown | - | Liss map (offspring) | - | SRR31053865 |
| **BW_0449** | $F_2$ *L.vul* × *L.mon* | Hatchling | unknown | - | Liss map (offspring) | - | SRR31053864 |
| **BW_0450** | $F_2$ *L.vul* × *L.mon* | Hatchling | unknown | - | Liss map (offspring) | - | SRR31053539 |
| **BW_0451** | $F_2$ *L.vul* × *L.mon* | Hatchling | unknown | - | Liss map (offspring) | - | SRR31053863 |
| **BW_0452** | $F_2$ *L.vul* × *L.mon* | Hatchling | unknown | - | Liss map (offspring) | - | SRR31053862 |
| **BW_0453** | $F_2$ *L.vul* × *L.mon* | Hatchling | unknown | - | Liss map (offspring) | - | SRR31053861 |
| **BW_0454** | $F_2$ *L.vul* × *L.mon* | Hatchling | unknown | - | Liss map (offspring) | - | SRR31053860 |
| **BW_0455** | $F_2$ *L.vul* × *L.mon* | Hatchling | unknown | - | Liss map (offspring) | - | SRR31053858 |
| **BW_0456** | $F_2$ *L.vul* × *L.mon* | Hatchling | unknown | - | Liss map (offspring) | - | SRR31053856 |
| **BW_0457** | $F_2$ *L.vul* × *L.mon* | Hatchling | unknown | - | Liss map (offspring) | - | SRR31053855 |
| **BW_0458** | $F_2$ *L.vul* × *L.mon* | Hatchling | unknown | - | Liss map (offspring) | - | SRR31053854 |
| **BW_0460** | $F_2$ *L.vul* × *L.mon* | Hatchling | unknown | - | Liss map (offspring) | - | SRR31053853 |
| **BW_0462** | $F_2$ *L.vul* × *L.mon* | Hatchling | unknown | - | Liss map (offspring) | - | SRR31053859 |
| **BW_0463** | $F_2$ *L.vul* × *L.mon* | Hatchling | unknown | - | Liss map (offspring) | - | SRR31053852 |
| **BW_0464** | $F_2$ *L.vul* × *L.mon* | Hatchling | unknown | - | Liss map (offspring) | - | SRR31053851 |
| **BW_0465** | $F_2$ *L.vul* × *L.mon* | Hatchling | unknown | - | Liss map (offspring) | - | SRR31053850 |
| **BW_0466** | $F_2$ *L.vul* × *L.mon* | Hatchling | unknown | - | Liss map (offspring) | - | SRR31053849 |
| **BW_0467** | $F_2$ *L.vul* × *L.mon* | Hatchling | unknown | - | Liss map (offspring) | - | SRR31053847 |

| | | | | | | | |
|---|---|---|---|---|---|---|---|
| **BW_0468** | $F_2$ *L.vul* × *L.mon* | Hatchling | unknown | - | Liss map (offspring) | - | SRR31053845 |
| **BW_0469** | $F_2$ *L.vul* × *L.mon* | Hatchling | unknown | - | Liss map (offspring) | - | SRR31053844 |
| **BW_0470** | $F_2$ *L.vul* × *L.mon* | Hatchling | unknown | - | Liss map (offspring) | - | SRR31053537 |
| **BW_0471** | $F_2$ *L.vul* × *L.mon* | Hatchling | unknown | - | Liss map (offspring) | - | SRR31053536 |
| **BW_0472** | $F_2$ *L.vul* × *L.mon* | Hatchling | unknown | - | Liss map (offspring) | - | SRR31053848 |
| **BW_0473** | $F_2$ *L.vul* × *L.mon* | Hatchling | unknown | - | Liss map (offspring) | - | SRR31053535 |
| **BW_0474** | $F_2$ *L.vul* × *L.mon* | Hatchling | unknown | - | Liss map (offspring) | - | SRR31053534 |
| **BW_0475** | $F_2$ *L.vul* × *L.mon* | Hatchling | unknown | - | Liss map (offspring) | - | SRR31053533 |
| **BW_0476** | $F_2$ *L.vul* × *L.mon* | Hatchling | unknown | - | Liss map (offspring) | - | SRR31053532 |
| **BW_0477** | $F_2$ *L.vul* × *L.mon* | Hatchling | unknown | - | Liss map (offspring) | - | SRR31053530 |
| **BW_0478** | $F_2$ *L.vul* × *L.mon* | Hatchling | unknown | - | Liss map (offspring) | - | SRR31053528 |
| **BW_0479** | $F_2$ *L.vul* × *L.mon* | Hatchling | unknown | - | Liss map (offspring) | - | SRR31053527 |
| **BW_0480** | $F_2$ *L.vul* × *L.mon* | Hatchling | unknown | - | Liss map (offspring) | - | SRR31053526 |
| **BW_0481** | $F_2$ *L.vul* × *L.mon* | Hatchling | unknown | - | Liss map (offspring) | - | SRR31053525 |
| **BW_0482** | $F_2$ *L.vul* × *L.mon* | Hatchling | unknown | - | Liss map (offspring) | - | SRR31053531 |
| **BW_0483** | $F_2$ *L.vul* × *L.mon* | Hatchling | unknown | - | Liss map (offspring) | - | SRR31053524 |
| **BW_0484** | $F_2$ *L.vul* × *L.mon* | Hatchling | unknown | - | Liss map (offspring) | - | SRR31053523 |
| **BW_0485** | $F_2$ *L.vul* × *L.mon* | Hatchling | unknown | - | Liss map (offspring) | - | SRR31053522 |
| **BW_0486** | $F_2$ *L.vul* × *L.mon* | Hatchling | unknown | - | Liss map (offspring) | - | SRR31053521 |
| **BW_0487** | $F_2$ *L.vul* × *L.mon* | Hatchling | unknown | - | Liss map (offspring) | - | SRR31053519 |
| **BW_0488** | $F_2$ *L.vul* × *L.mon* | Hatchling | unknown | - | Liss map (offspring) | - | SRR31053517 |
| **BW_0489** | $F_2$ *L.vul* × *L.mon* | Hatchling | unknown | - | Liss map (offspring) | - | SRR31053516 |
| **BW_0490** | $F_2$ *L.vul* × *L.mon* | Hatchling | unknown | - | Liss map (offspring) | - | SRR31053843 |
| **BW_0491** | $F_2$ *L.vul* × *L.mon* | Hatchling | unknown | - | Liss map (offspring) | - | SRR31053842 |
| **BW_0492** | $F_2$ *L.vul* × *L.mon* | Hatchling | unknown | - | Liss map (offspring) | - | SRR31053520 |
| **BW_0493** | $F_2$ *L.vul* × *L.mon* | Hatchling | unknown | - | Liss map (offspring) | - | SRR31053841 |
| **BW_0494** | $F_2$ *L.vul* × *L.mon* | Hatchling | unknown | - | Liss map (offspring) | - | SRR31053840 |
| **BW_0495** | $F_2$ *L.vul* × *L.mon* | Hatchling | unknown | - | Liss map (offspring) | - | SRR31053839 |
| **BW_0496** | $F_2$ *L.vul* × *L.mon* | Hatchling | unknown | - | Liss map (offspring) | - | SRR31053838 |
| **BW_0497** | $F_2$ *L.vul* × *L.mon* | Hatchling | unknown | - | Liss map (offspring) | - | SRR31053836 |
| **BW_0498** | $F_2$ *L.vul* × *L.mon* | Hatchling | unknown | - | Liss map (offspring) | - | SRR31053834 |
| **BW_0499** | $F_2$ *L.vul* × *L.mon* | Hatchling | unknown | - | Liss map (offspring) | - | SRR31053833 |
| **BW_0500** | $F_2$ *L.vul* × *L.mon* | Hatchling | unknown | - | Liss map (offspring) | - | SRR31053832 |
| **BW_0501** | $F_2$ *L.vul* × *L.mon* | Hatchling | unknown | - | Liss map (offspring) | - | SRR31053831 |
| **BW_0502** | $F_2$ *L.vul* × *L.mon* | Hatchling | unknown | - | Liss map (offspring) | - | SRR31053830 |
| **BW_0503** | $F_2$ *L.vul* × *L.mon* | Hatchling | unknown | - | Liss map (offspring) | - | SRR31053829 |
| **BW_0504** | $F_2$ *L.vul* × *L.mon* | Hatchling | unknown | - | Liss map (offspring) | - | SRR31053828 |
| **BW_0505** | $F_2$ *L.vul* × *L.mon* | Hatchling | unknown | - | Liss map (offspring) | - | SRR31053827 |
| **BW_0506** | $F_2$ *L.vul* × *L.mon* | Hatchling | unknown | - | Liss map (offspring) | - | SRR31053826 |
| **BW_0507** | $F_2$ *L.vul* × *L.mon* | Hatchling | unknown | - | Liss map (offspring) | - | SRR31053825 |
| **BW_0508** | $F_2$ *L.vul* × *L.mon* | Hatchling | unknown | - | Liss map (offspring) | - | SRR31053823 |
| **BW_0510** | $F_2$ *L.vul* × *L.mon* | Hatchling | unknown | - | Liss map (offspring) | - | SRR31053822 |
| **BW_0511** | $F_2$ *L.vul* × *L.mon* | Hatchling | unknown | - | Liss map (offspring) | - | SRR31053821 |
| **BW_0512** | $F_2$ *L.vul* × *L.mon* | Hatchling | unknown | - | Liss map (offspring) | - | SRR31053625 |
| **BW_0513** | $F_2$ *L.vul* × *L.mon* | Hatchling | unknown | - | Liss map (offspring) | - | SRR31053624 |
| **BW_0514** | $F_2$ *L.vul* × *L.mon* | Hatchling | unknown | - | Liss map (offspring) | - | SRR31053837 |
| **BW_0515** | $F_2$ *L.vul* × *L.mon* | Hatchling | unknown | - | Liss map (offspring) | - | SRR31053623 |
| **BW_0516** | $F_2$ *L.vul* × *L.mon* | Hatchling | unknown | - | Liss map (offspring) | - | SRR31053622 |
| **BW_0517** | $F_2$ *L.vul* × *L.mon* | Hatchling | unknown | - | Liss map (offspring) | - | SRR31053621 |
| **BW_0518** | $F_2$ *L.vul* × *L.mon* | Hatchling | unknown | - | Liss map (offspring) | - | SRR31053620 |
| **BW_0519** | $F_2$ *L.vul* × *L.mon* | Hatchling | unknown | - | Liss map (offspring) | - | SRR31053617 |
| **BW_0520** | $F_2$ *L.vul* × *L.mon* | Hatchling | unknown | - | Liss map (offspring) | - | SRR31053616 |
| **BW_0521** | $F_2$ *L.vul* × *L.mon* | Hatchling | unknown | - | Liss map (offspring) | - | SRR31053615 |
| **BW_0522** | $F_2$ *L.vul* × *L.mon* | Hatchling | unknown | - | Liss map (offspring) | - | SRR31053614 |
| **BW_0523** | $F_2$ *L.vul* × *L.mon* | Hatchling | unknown | - | Liss map (offspring) | - | SRR31053613 |
| **BW_0524** | $F_2$ *L.vul* × *L.mon* | Hatchling | unknown | - | Liss map (offspring) | - | SRR31053612 |

| | | | | | | | |
|---|---|---|---|---|---|---|---|
| **BW_0525** | $F_2$ *L.vul* × *L.mon* | Hatchling | unknown | - | Liss map (offspring) | - | SRR31053611 |
| **BW_0526** | $F_2$ *L.vul* × *L.mon* | Hatchling | unknown | - | Liss map (offspring) | - | SRR31053610 |
| **BW_0527** | $F_2$ *L.vul* × *L.mon* | Hatchling | unknown | - | Liss map (offspring) | - | SRR31053609 |
| **BW_0528** | $F_2$ *L.vul* × *L.mon* | Hatchling | unknown | - | Liss map (offspring) | - | SRR31053608 |
| **BW_0529** | $F_2$ *L.vul* × *L.mon* | Hatchling | unknown | - | Liss map (offspring) | - | SRR31053606 |
| **BW_0530** | $F_2$ *L.vul* × *L.mon* | Hatchling | unknown | - | Liss map (offspring) | - | SRR31053602 |
| **BW_0531** | $F_2$ *L.vul* × *L.mon* | Hatchling | unknown | - | Liss map (offspring) | - | SRR31053603 |
| **BW_0532** | $F_2$ *L.vul* × *L.mon* | Hatchling | unknown | - | Liss map (offspring) | - | SRR31053737 |
| **BW_0533** | $F_2$ *L.vul* × *L.mon* | Hatchling | unknown | - | Liss map (offspring) | - | SRR31053736 |
| **BW_0534** | $F_2$ *L.vul* × *L.mon* | Hatchling | unknown | - | Liss map (offspring) | - | SRR31053604 |
| **BW_0535** | $F_2$ *L.vul* × *L.mon* | Hatchling | unknown | - | Liss map (offspring) | - | SRR31053735 |
